# The RBR E3 ubiquitin ligase HOIL-1 can ubiquitinate diverse non-protein substrates *in vitro*

**DOI:** 10.1101/2024.11.29.626046

**Authors:** Xiangyi S. Wang, Jenny Jiou, Anthony Cerra, Simon A. Cobbold, Marco Jochem, Ka Hin Toby Mak, Leo Corcilius, Richard J. Payne, Ethan D. Goddard-Borger, David Komander, Bernhard C. Lechtenberg

## Abstract

HOIL-1 is a RING-between-RING (RBR)-family E3 ubiquitin ligase and component of the linear ubiquitin chain assembly complex (LUBAC). While most E3 ubiquitin ligases conjugate ubiquitin to protein lysine sidechains, HOIL-1 has been reported to ubiquitinate hydroxyl groups in protein serine and threonine sidechains and glucosaccharides, such as glycogen and its building block maltose, *in vitro*. However, HOIL-1 substrate specificity is currently poorly defined. Here we show that HOIL-1 is unable to ubiquitinate lysine but can efficiently ubiquitinate serine as well as a variety of model and physiologically relevant di- and monosaccharides *in vitro*. We identify a critical catalytic histidine residue, His510, in the flexible catalytic site of HOIL-1 that enables this O-linked ubiquitination and prohibits ubiquitin discharge onto lysine sidechains. Finally, we utilise HOIL-1’s *in vitro* non-proteinaceous ubiquitination activity and an engineered, constitutively active HOIL-1 variant to produce preparative amounts of different ubiquitinated saccharides that can be used as tool compounds and standards in the rapidly emerging field of non-proteinaceous ubiquitination.

## INTRODUCTION

Ubiquitination is a post-translational modification characterised by the conjugation of the small, 76-amino acid protein ubiquitin to other proteins. Ubiquitination is mediated by a three-tier cascade of enzymes, the E1 ubiquitin activating enzyme, the E2 ubiquitin conjugating enzymes and the E3 ubiquitin ligases (1). Ubiquitin is commonly defined as a post-translational protein modification where the ubiquitin C-terminus forms an isopeptide bond with a substrate lysine sidechain (2). Ubiquitin itself contains seven internal lysine residues, allowing ubiquitin to form different types of polyubiquitin chains, each with different biological functions (2). Ubiquitin can also form peptide bonds with the N-terminal amine of proteins, including ubiquitin itself (2). In humans, these M1-linked (or linear) ubiquitin chains are thought to be exclusively formed by RING-between-RING (RBR) E3 ubiquitin ligase HOIP, the catalytic centre of the linear ubiquitin chain assembly complex (LUBAC) (3–9).

In addition to the canonical ubiquitination on amines, non-canonical protein ubiquitination on the hydroxyl groups of Ser and Thr residues via oxyester bonds (O-linked ubiquitination) or the sulfhydryl group of Cys residues via thioester bonds have also been described (10–12). Over the past years, researchers have identified multiple E3 ubiquitin ligases that can form ubiquitin oxyester linkages. MYCBP2 utilises a specific Cys-relay system to transfer ubiquitin to Thr residues (13). The RBR E3 ligase HHARI (ARIH1) can ubiquitinate Ser or Thr residues on cyclin E and Nrf1 (14, 15). The RBR ligase and LUBAC component HOIL-1, which previously showed enigmatic E3 ligase activity (8, 16), has recently been suggested to ubiquitinate Ser and Thr residues on other LUBAC components, components of myddosomes and ubiquitin (17–20).

In addition to ubiquitin as a protein post-translational modification, since 2021, the array of potential ubiquitin substrates has been expanded to non-proteinaceous biomolecules. Ubiquitin has been found conjugated to bacterial lipopolysaccharides (LPS), e.g. in Salmonella-infected cells (21), lipids (22), ADP-ribose attached to proteins or nucleic acids (23, 24), saccharides (15, 25), single stranded RNA and DNA (26) and synthetic small molecules (27). One of the E3 ligases implicated in ubiquitination of glucosaccharides is HOIL-1, which can ubiquitinate glycogen, the main storage form of glucose in humans, and its building block maltose *in vitro* (25, 28–30). The diverse set of reported HOIL-1 activities targeting Lys and Ser/Thr sidechains and glucosaccharides prompted us to investigate the relative activity of HOIL-1 against different classes of reported HOIL-1 substrates in a more quantitative approach.

Progress in the ubiquitin field over the past two decades was largely driven by the development of highly specific tools and technologies, ranging from chain-type specific antibodies and affimers (31–34), to antibody-based enrichment strategies (35–37) and selective ubiquitin proteomics (38–40). However, many of these tools are only suitable for Lys- or protein-conjugated ubiquitin. Therefore, while non-proteinaceous ubiquitination is a rapidly emerging field, progress is currently hampered by the lack of tools and standards to study these novel modifications (41).

Here, we characterise the *in vitro* substrate specificity of recombinant human HOIL-1, showing that HOIL-1 can efficiently ubiquitinate Ser residues and different saccharides, with weak activity for Thr and no activity for Lys residues. We identify HOIL-1 His510 as a key active site residue that discriminates between ubiquitination of hydroxyl groups in Ser/Thr residues and ε-amine groups in Lys residues. Our work shows that HOIL-1 can ubiquitinate a broad range of saccharides tested with only small differences in the relative activity. We utilise human HOIL-1 and an engineered, constitutively active HOIL-1 variant to efficiently generate preparative, milligram amounts of various highly pure ubiquitinated saccharides. These ubiquitinated saccharides will be essential as novel tool compounds and standards to investigate non-proteinaceous ubiquitination by HOIL-1 and other E3 ligases and to test recognition of these ubiquitin signals by specific receptors and their removal by deubiquitinases (DUBs). To this end, in a proof-of-concept experiment, we show that several human DUBs can cleave ubiquitinated maltose *in vitro*. Together, our work provides new insights into HOIL-1 *in vitro* activity and specificity and an avenue to develop novel tools to investigate non-proteinaceous ubiquitination.

## RESULTS

### HOIL-1 forms O-linked ubiquitin on Ser/Thr-containing peptides

To compare HOIL-1 E3 ligase activity for different amino acid acceptors, we reconstituted HOIL-1 E3 ligase activity *in vitro* using recombinant E1, the E2 UbcH7 and full-length human HOIL-1 in the presence of M1-linked di-ubiquitin, which was previously shown to allosterically activate HOIL-1 (25, 28–30). We compare HOIL-1 activity against different pentameric peptides with the general sequence Ac-EGxGN-NH_2_, where the central amino acid x is either a Lys (K), Ser (S), Thr (T) or Arg (R) residue (13). Lys contains an amine as an acceptor, Ser and Thr contain a hydroxyl group acceptor, whereas the Arg sidechain (bearing a guanidine functional group in the side chain) is not expected to be an acceptor for ubiquitination.

Our experiments show that HOIL-1 can efficiently attach ubiquitin to the Ser-containing peptide as indicated by disappearance of the ubiquitin substrate band and appearance of a slower migrating ubiquitin-peptide band via SDS-PAGE (Fig 1A). Under our reaction conditions, about half of the ubiquitin is conjugated to the Ser-containing peptide in ∼45 min (Fig 1A, B). We also observe formation of a ubiquitinated peptide with the Thr-containing peptide, albeit to a lower degree with only about 20% of ubiquitin conjugated to the peptide after 45 min (Fig 1A, B). In contrast to the activity with the hydroxyl group-containing Ser and Thr peptides, we do not observe ubiquitination of the Lys- and Arg-containing peptides with WT HOIL-1, even though HOIL-1 in this assay is clearly active, as indicated by HOIL-1 auto-ubiquitination (Fig 1C).

**Figure 1:**
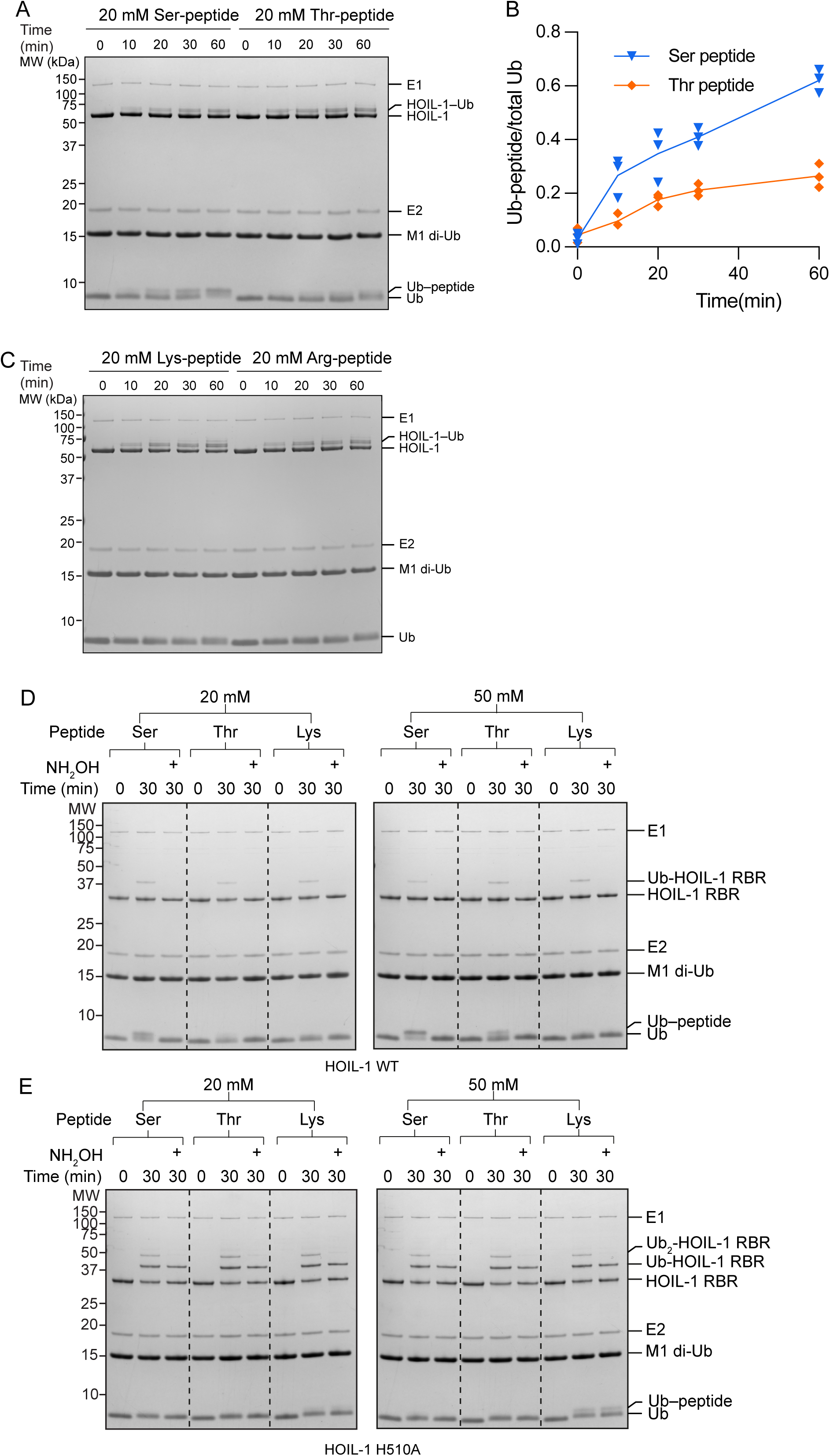
HOIL-1 ubiquitination of Ser and Thr residues *in vitro* is dependent on His510. **(A)** Peptide ubiquitination assay with 20 mM synthetic Ser- or Thr-containing peptides (Ac-EGxGN-NH_2_, where is x is either Ser or Thr). **(B)** Quantification of HOIL-1-mediated peptide ubiquitination reactions performed as represented in panel A. The proportion of ubiquitinated peptide over total ubiquitin is plotted over time. Individual data points and connecting lines of the mean are shown (n=3). **(C)** Peptide ubiquitination assays as in panel A with Lys- or Arg-containing peptides. This experiment was performed three times with consistent results. **(D)** Peptide ubiquitination assay with 20 mM (left) and 50 mM (right) synthetic Ser-, Thr-, or Lys-containing peptides and WT HOIL-1. Samples were taken at the indicated time points and analysed via SDS-PAGE under reducing conditions. Half of the samples taken at 30 min were treated with hydroxylamine (NH_2_OH) as indicated. **(E)** Peptide ubiquitination assay as in panel D but with the HOIL-1 H510A mutant. Experiments in panels D and E were performed at least twice (except for the right gel in panel E) with consistent results. Representative gels are shown. Ub: ubiquitin.

### HOIL-1 His510 enables O-linked ubiquitination and prohibits ubiquitin discharge onto Lys residues

HOIL-1’s C-terminal residue, His510, forms part of HOIL-1’s catalytic triad and is critical for maltose ubiquitination (28–30). RBR E3 ubiquitin ligases like HOIL-1 catalyse ubiquitination in a two-step mechanism where ubiquitin is first transferred from the E2 active site Cys to the E3 active site Cys before being transferred to the substrate (42, 43). We previously showed that mutation of His510 to Ala (H510A) does not affect E2-ubiquitin discharge (29), indicating that His510 is critical only for the second catalytic step. Interestingly, while a HOIL-1 H510A mutant was inactive in ubiquitinating maltose, we previously observed enhanced auto-ubiquitination of the HOIL-1 H510A mutant compared to WT HOIL-1 (29). These observations suggest that HOIL-1 His510 plays a complex role in ubiquitin transfer from the HOIL-1 active site to different substrates.

To further investigate the altered activity of the HOIL-1 H510A mutant and compare the role of His510 in ubiquitinating different acceptors, we compared HOIL-1 WT and H510A activity in the peptide ubiquitination assays. As before, WT HOIL-1 efficiently ubiquitinated the Ser-peptide and, to a lesser extent, the Thr-peptide (Fig 1D). The slower migrating band disappears upon treatment with hydroxylamine (NH_2_OH) which specifically breaks oxyester bonds while leaving (iso)-peptide bonds intact (13), supporting that the peptides are ubiquitinated on the Ser/Thr sidechains. In contrast to WT HOIL-1, the HOIL-1 H510A mutant does not modify the Ser/Thr peptides even when using a higher substrate concentration (Fig 1E). However, we observe strong autoubiquitination with HOIL-1 H510A compared to WT HOIL-1 (Fig 1D,E). Surprisingly, while the HOIL-1 WT autoubiquitination disappears after treatment with hydroxylamine (Fig 1D), most of the HOIL-1 H510A auto-ubiquitination is resistant to hydroxylamine treatment when assayed on SDS-PAGE under reducing conditions (Fig 1E), demonstrating that the autoubiquitination is via isopeptide-linked ubiquitination on a Lys sidechain, rather than Ser/Thr ubiquitination or ubiquitin loaded to the catalytic Cys. We therefore investigated the activity of HOIL-1 WT and H510A with the Lys-peptide. As before, we do not observe Lys-peptide ubiquitination by WT HOIL-1 (Fig 1D). However, using HOIL-1 H510A, we detect a faint modification of the Lys-peptide with 20 mM substrate, with a more pronounced product formation with 50 mM substrate (Fig 1E). These modifications are hydroxylamine-resistant, indicating that they are isopeptide-linked via the lysine sidechain, in line with our observations of HOIL-1 auto-ubiquitination.

Together, these results confirm that HOIL-1 His510 is required for oxyester-linked ubiquitination, but at the same time prevents ubiquitination of Lys sidechains. His510 therefore is a critical switch to fine-tune HOIL-1 activity towards oxyester linked ubiquitination.

### Ubiquitin binding modulates the HOIL-1 active site conformation

To investigate the importance of His510 for HOIL-1 catalytic activity, we determined the structure of the HOIL-1 RING2 catalytic domain with the active site C460A mutation in the presence of ubiquitin (Fig 2, Fig EV1, Table 1). The asymmetric unit of our structure contains two copies of the HOIL-1 RING2 domain, one of which is loaded with ubiquitin (Fig 2A, Fig EV1A). The overall fold of the HOIL-1 RING2 domain is the same as in previously determined HOIL-1 RING2 structures, with an α-helix preceding the RING2 domain and the HOIL-1-exclusive 2Zn/6Cys binuclear cluster forming the C-terminal portion of the RING2 (Fig EV1B) (28–30). The ubiquitin Ile44 patch interacts with the HOIL-1 helix, with the ubiquitin C-terminus pointing into the HOIL-1 active site (Fig 2B). The ubiquitin therefore mimics the donor ubiquitin as observed in the E2-ubiquitin/HOIL-1 transthiolation complex (Fig EV1B) (29).

**Figure 2:**
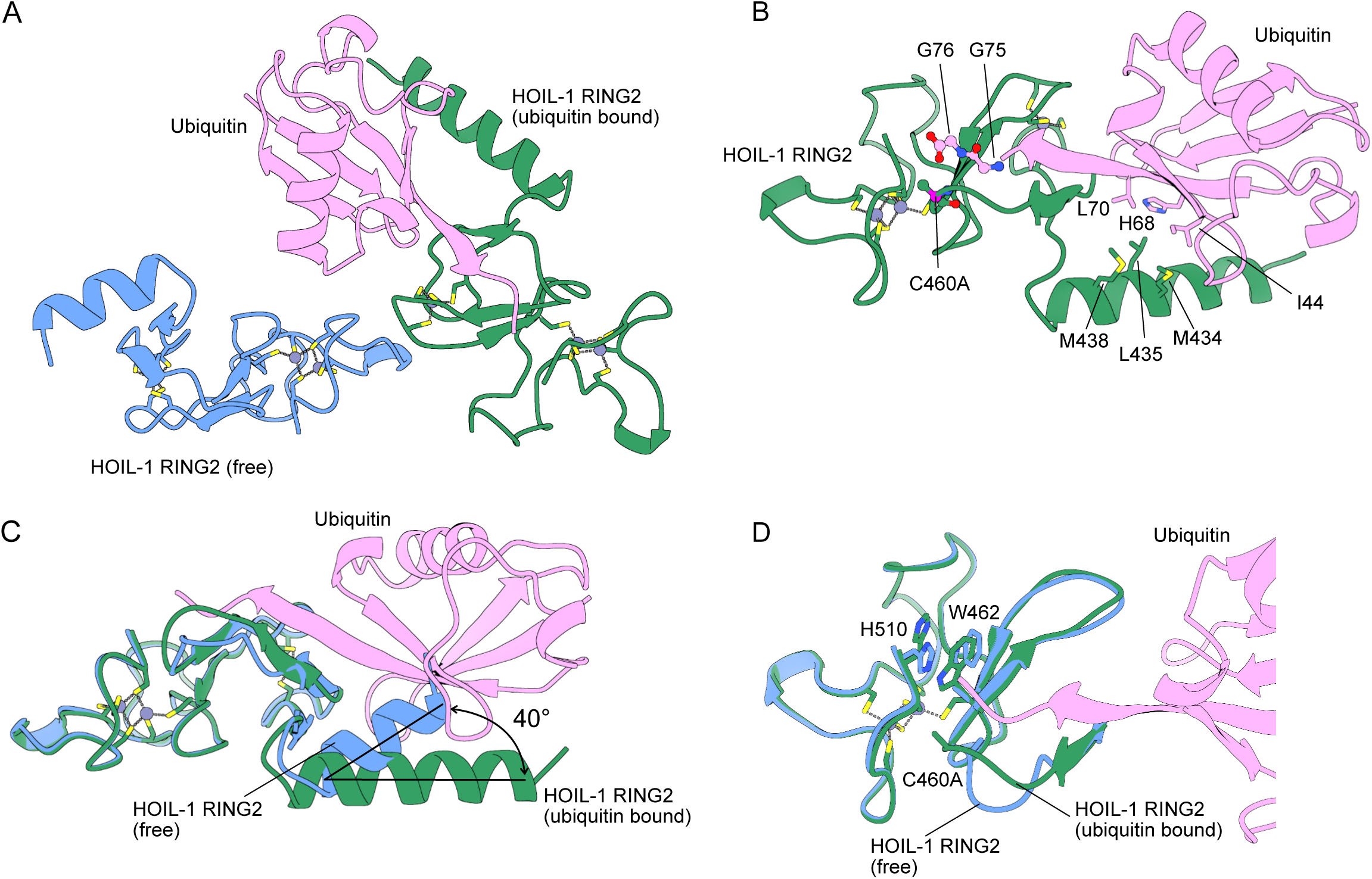
Crystal structure of HOIL-1 RING2 domain bound by ubiquitin. **(A)** Overview of the crystallographic asymmetric unit shows two HOIL-1 RING2 domains (green, blue) with one of them (green) bound to ubiquitin (pink). Bound zinc ions are shown as grey spheres with zinc ion-coordinating amino acid sidechains shown as sticks. **(B)** Detailed view of the interactions between ubiquitin (pink) and the HOIL-1 RING2 domain (green). Key interacting residues are shown as sticks and labelled. **(C)** Structural superposition of the HOIL-1 RING2 domains in the free state (blue) and bound to ubiquitin (green with ubiquitin in pink) highlight the conformational change between the core RING2 domain and the N-terminal helix upon ubiquitin binding. **(D)** Structural superposition of the HOIL-RING2 domains in the free state (blue) and bound to ubiquitin (green with ubiquitin in pink) highlight the conformational changes in the active site loop, W462 and H510. The active site Cys460 in the structure is mutated to Ala (C460A). The N-terminal part of the HOIL-1 RING2 domain is omitted for clarity.

**Table 1:**
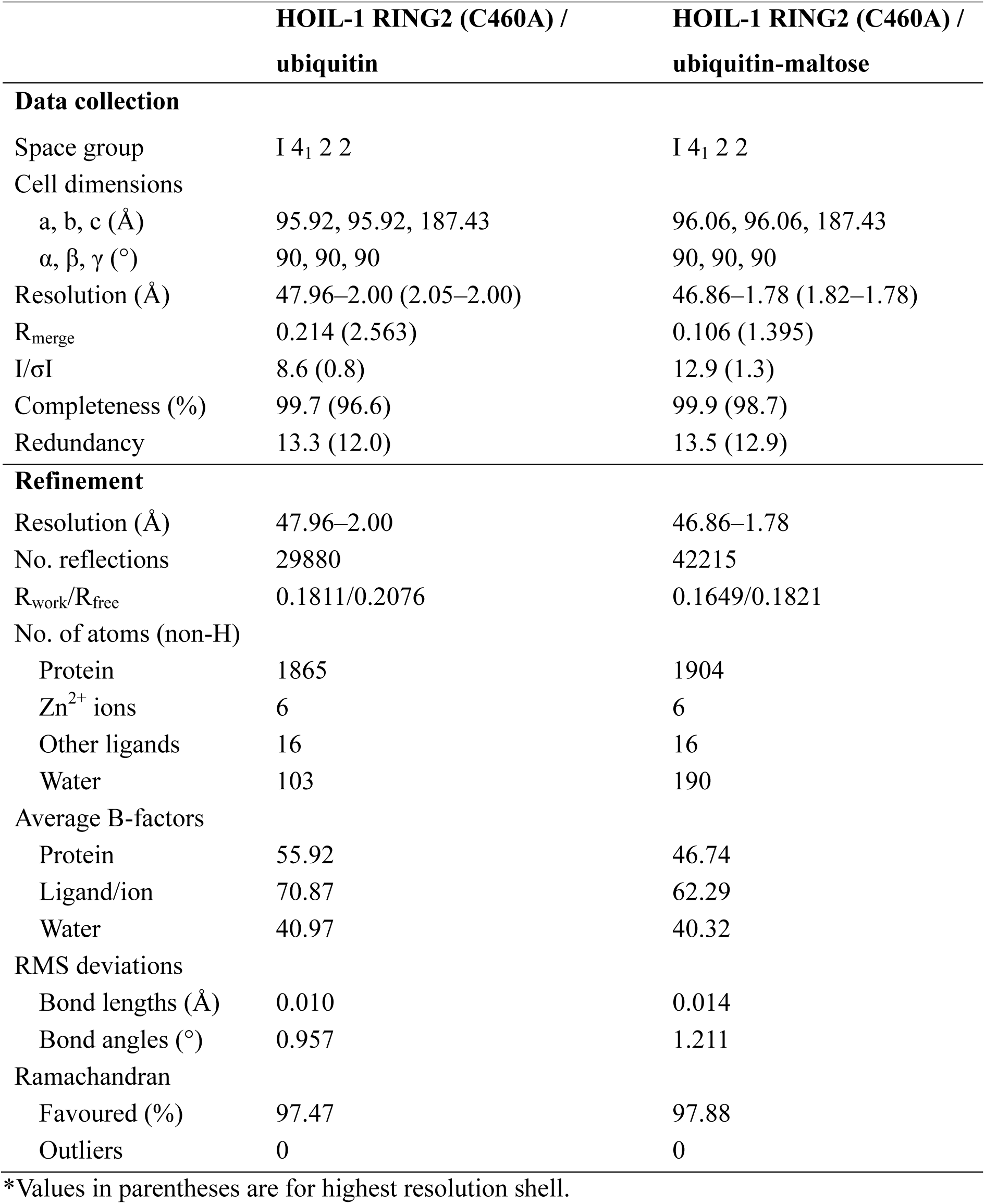
Crystallographic data collection, processing and refinement statistics.

Comparison of the ubiquitin-bound HOIL-1 RING2 to the unbound RING2 in the asymmetric unit enables us to investigate the conformational changes induced by ubiquitin binding. Ubiquitin-binding does not globally alter the HOIL-1 RING2 fold, but we observe two distinct changes. First, ubiquitin-binding alters the angle between the RING2 domain and its N-terminal helix by 30–40° (Fig 2C, Fig EV1C). This observation supports the notion that RBR ligases are highly dynamic enzymes that undergo vast conformational changes throughout their catalytic cycle to allow for binding of their different interaction partners (29, 43, 44). Notably, the conformational changes are not restricted to the N-terminal helix, but ubiquitin-binding also rearranges the HOIL-1 active site. The biggest differences are observed in the protein backbone around the active site Cys460 residue (mutated to Ala in the structure), between Gln455 and Asp461, and in the sidechain orientations of Trp462 and His510 (Fig 2D). In the ubiquitin-bound structure, Gln455 – Lys457 form an extended β-sheet with the ubiquitin C-terminus, whereas the same region in the unbound HOIL-1 RING2 adopts a less structured conformation (Fig 2D). The ubiquitin C-terminus also leads to a different rotamer of the Trp462 sidechain, where Trp462 is flipped towards the active site Cys460, leading to re-arrangement of the His510 sidechain away from the active site (Fig 2D). These re-arrangements in the active site upon ubiquitin binding closely match the conformation previously observed in the HOIL-1/E2-ubiquitin/ubiquitin thioester transfer complex structure (Fig EV1D) (29), suggesting that the observed conformation presents the catalytically active RING2 conformation.

### HOIL-1 prefers maltose over Ser/Thr containing peptides as substrates

As outlined before, HOIL-1 has been reported to ubiquitinate hydroxyl groups in Ser/Thr residues and other biomolecules, such as glucosaccharides. Therefore, after establishing that HOIL-1 efficiently ubiquitinates hydroxyl groups in Ser-residues over Lys residues, we investigated if HOIL-1 prefers hydroxyl groups in amino acids over hydroxyl groups in oligosaccharides. Our data show that HOIL-1 can ubiquitinate maltose more efficiently than the Ser- or Thr-containing peptides at the same substrate concentrations (Fig 3A, Fig EV2A). Notably, even though maltose is a disaccharide consisting of two α-1,4-linked glucose molecules (Fig 3B), we do not observe any evidence of dual ubiquitinated maltose species in our gel-based assays or matrix-assisted laser desorption/ionisation time-of-flight (MALDI-TOF) mass spectrometry (Fig EV2A,B) in line with previous work showing that the larger oligosaccharides maltoheptaose and α-cyclodextrin are mono-ubiquitinated on the C6 hydroxyl group of only one of their glucose subunits (25, 30).

**Figure 3:**
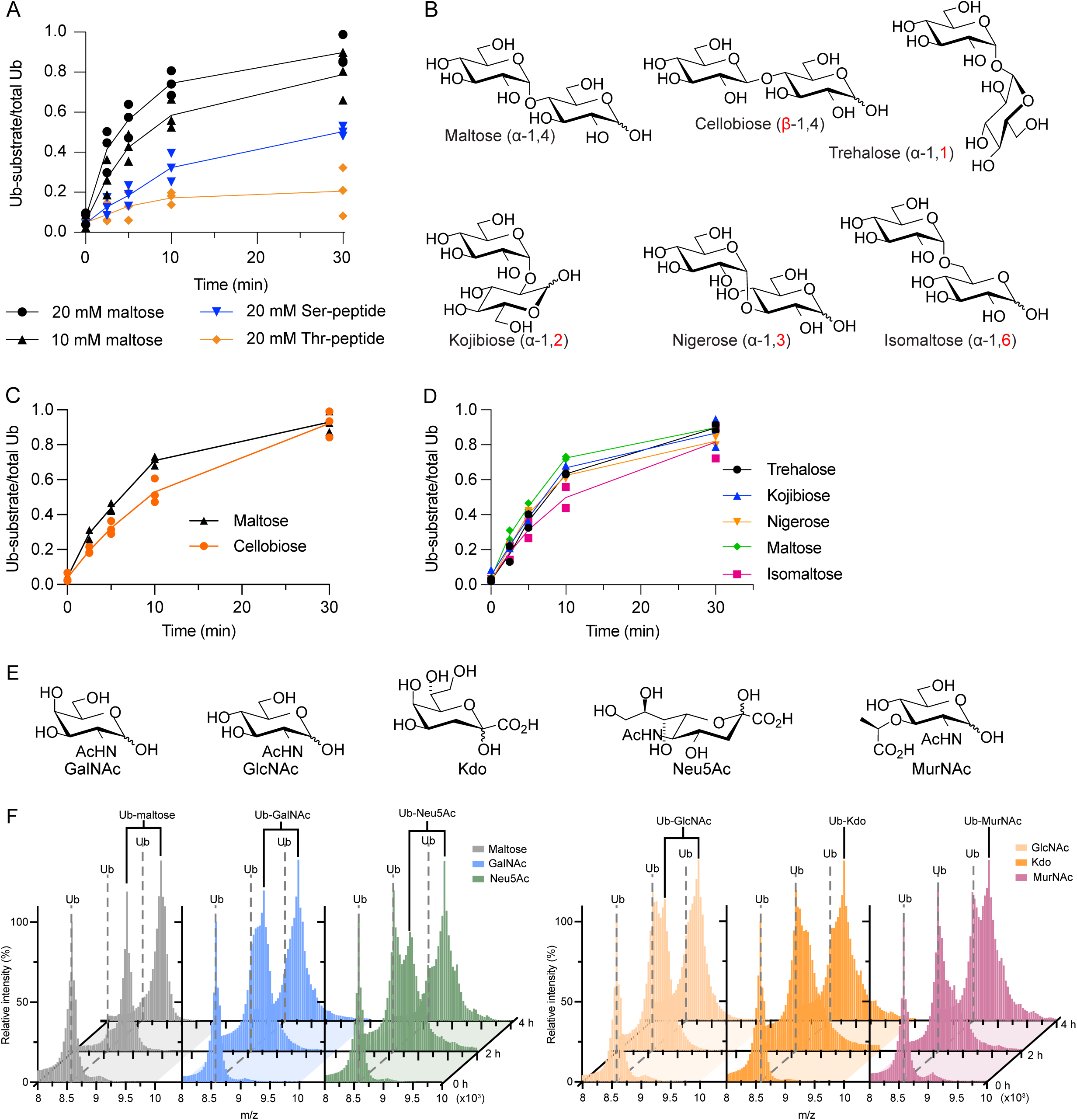
HOIL-1 ubiquitinates multiple disaccharides and physiological monosaccharides *in vitro*. **(A)** Quantification of HOIL-1-mediated *in vitro* substrate ubiquitination comparing Ser- and Thr-peptides with maltose. The proportion of ubiquitinated substrate over total ubiquitin as quantified from Coomassie-stained SDS-PAGE gels is plotted over time. Individual data points and connecting lines of the mean are shown (n=3). **(B)** Chemical structures of disaccharide substrates tested. Differences to maltose are highlighted in red font. **(C)** Quantification of HOIL-1-mediated *in vitro* substrate ubiquitination reactions comparing the disaccharides maltose and cellobiose. Individual data points and connecting lines of the mean are shown (n=3). **(D)** Quantification of HOIL-1-mediated *in vitro* substrate ubiquitination reactions comparing disaccharides as indicated. Individual data points and connecting lines of the mean are shown (n=3). Data in panels C and D are from the same experiment. **(E)** Chemical structures of the physiological monosaccharide substrates tested. **(F)** MALDI-TOF analysis of HOIL-1-mediated *in vitro* substrate ubiquitination reactions of the substrates shown in panel E. Experiments were performed twice with consistent results. Exemplary data are shown. Ub: ubiquitin.

### HOIL-1 ubiquitinates different model disaccharides with little specificity

There is a great deal of diversity in oligosaccharide structure due to their complex biochemistry that allows for different linkages of two D-glucose molecules to form ten disaccharide isomers, including α- and β-linkages, as in maltose (α-1,4) and cellobiose (β-1,4), and linkages between different positions of each sugar unit (regioselectivity): α-1,1 in trehalose, α-1,2 in kojibiose, α-1,3 in nigerose, α-1,4 in maltose and α-1,6 in isomaltose (Fig 3B). While many of these disaccharides are not produced in humans, they enable systematic interrogation of HOIL-1 substrate specificity, or lack thereof. A comparison of maltose (α-1,4) and cellobiose (β-1,4) shows that HOIL-1 can ubiquitinate both disaccharides to a similar level after 30 min, but maltose shows a slightly higher initial rate of ubiquitination, suggesting that maltose may be a preferred substrate for HOIL-1, although the difference in the initial rate of transfer (up to 10 min) is only about 1.4-fold (Fig 3C, Fig EV2C).

Next, we compared different disaccharides with regioselectivity. We restricted our analysis to α glycosidic linkages as found in the preferred substrate maltose. As before, we only observe small differences in HOIL-1 activity using maltose or any of the other disaccharides as substrates (Fig 3D, Fig EV2C). The biggest difference, 1.4-fold, was observed between maltose (α-1,4) and isomaltose (α-1,6) (Fig 3D). In summary, HOIL-1 can ubiquitinate all di-saccharides tested with only marginal differences in the rate of ubiquitination, suggesting that HOIL-1 has broad substrate specificity for different model di-saccharides.

### HOIL-1 ubiquitinates a variety of physiological monosaccharides *in vitro*

So far, we systematically investigated HOIL-1 activity on different model disaccharides including many with limited physiological relevance in humans. To investigate more physiologically relevant potential HOIL-1 substrates, we selected multiple saccharides that may be present in a cellular environment in humans (Fig 3E). These include *N*-acetyl-D-galactosamine (GalNAc) and *N*-acetyl-D-glucosamine (GlcNAc), which are commons sugars found in O-linked protein glycosylation in humans; ketodeoxyoctonic acid (Kdo) and *N*-acetylmuramic acid (MurNAc) found on the surface of bacterial pathogens that may infect human cells; and *N*-acetylneuraminic acid (Neu5Ac), a sialic acid that is a common terminal motif on mammalian glycoconjugates. We used the same *in vitro* assay as before, but instead of trying to resolve and detect products by SDS-PAGE, we used MALDI-TOF MS to distinguish between modified and unmodified ubiquitin species. To simplify analysis via MALDI-TOF, we omitted the allosteric activator M1-linked di-ubiquitin and instead increased the reaction time from 0.5 to 4 h.

Our control experiment with maltose confirms that HOIL-1 can ubiquitinate maltose under these conditions and shows that MALDI-TOF MS can efficiently discriminate between free ubiquitin and ubiquitinated maltose (Fig 3F). Likewise, HOIL-1 ubiquitinates GalNAc, Neu5Ac, GlcNAc, Kdo and MurNAc within 4 h (Fig 3F), albeit with different efficiency. While maltose is fully modified, only about 50% of the ubiquitin is conjugated to MurNAc, suggesting that MurNAc ubiquitination by HOIL-1 is less efficient than maltose ubiquitination. The 2 h-timepoint reveals more subtle differences between the different substrates with near complete conversion of ubiquitin to ubiquitin-saccharide with maltose and GalNAc (Fig 3F). We only observe 50% conversion with Neu5Ac, GlcNAc and Kdo, whereas less than half of the ubiquitin is conjugated to MurNAc (Fig 3F). Together, these experiments show that HOIL-1 can efficiently ubiquitinate a variety of mono- and di-saccharides *in vitro*.

### HOIL-1-mediated production of preparative amounts of ubiquitinated saccharides

Ubiquitination of non-proteinaceous substrates, such as LPS, lipids and saccharides, is of growing interest in the ubiquitin field (10–12, 21–26, 30, 45). However, tools to study these novel modifications are currently very limited (41). Therefore, the scaled-up chemoenzymatic preparation of ubiquitinated non-proteinaceous substrates could provide standards to benchmark new quantitative methods or probes to identify and characterise novel ubiquitin-substrate binding domains or DUBs. As we observe that HOIL-1 can efficiently and specifically ubiquitinate a wide variety of saccharides, we hypothesised that HOIL-1 would be an ideal enzyme to generate preparative quantities of ubiquitinated non-proteinaceous substrates.

To this end, we modified our *in vitro* HOIL-1 ubiquitination assay to achieve higher yields of ubiquitinated product and reaction completion to enable purification of the ubiquitinated product. As a proof of principle, we used maltose as substrate, since we had determined it to be the preferred substrate *in vitro*. The key modifications were to use a ten-fold higher concentration of ubiquitin (100 µM) and to increase the reaction time to up to 16 h. We also used an N-terminally His-tagged ubiquitin to enable purification of the ubiquitinated maltose from unreacted maltose and other reaction components.

Comparing the reaction rate of HOIL-1 in a small-scale time-course experiment using 10 µM or 100 µM His-tagged ubiquitin shows that HOIL-1 efficiently modifies 50% of 10 µM ubiquitin within 2 h with full conversion to ubiquitin-maltose after 16 h (Fig 4A). Under the same enzyme concentrations but with 100 µM ubiquitin, we observe that after 2 h, only about 25% of the ubiquitin is conjugated to maltose (Fig 4A). Incubation overnight (∼16 h) results in full ubiquitin conversion to the slower migrating band without evidence of secondary reaction products, for example ubiquitin chains (Fig 4A). Based on these promising results, we developed a protocol to generate and purify milligram quantities of ubiquitin-maltose (Fig 4B–E). We scaled up the reaction to a 5 ml volume and used 100 µM His-tagged ubiquitin (∼1 mg/ml) for a total input of 5 mg ubiquitin. Reaction for 16 h resulted in complete conversion to the slower migrating His-tagged ubiquitin-maltose band (Fig 4C). Subsequent purification via Ni-NTA affinity chromatography resulted in enrichment of His-tagged ubiquitin-maltose (Fig 4C). Final purification by size-exclusion chromatography yielded His-tagged ubiquitin-maltose at excellent purity as determined by SDS-PAGE (Fig 4D) and native mass spectrometry (Fig 4E). Using this protocol, we can typically generate 2.5 mg highly pure ubiquitin-maltose from 5 mg ubiquitin input. Our method is applicable to a wide variety of saccharides including GalNAc, GlcNAc, N-acetyllactosamine (LacNAc) and mucin-type O-glycans Tn, T and sTn antigens α-linked to serine (Fig EV3).

**Figure 4:**
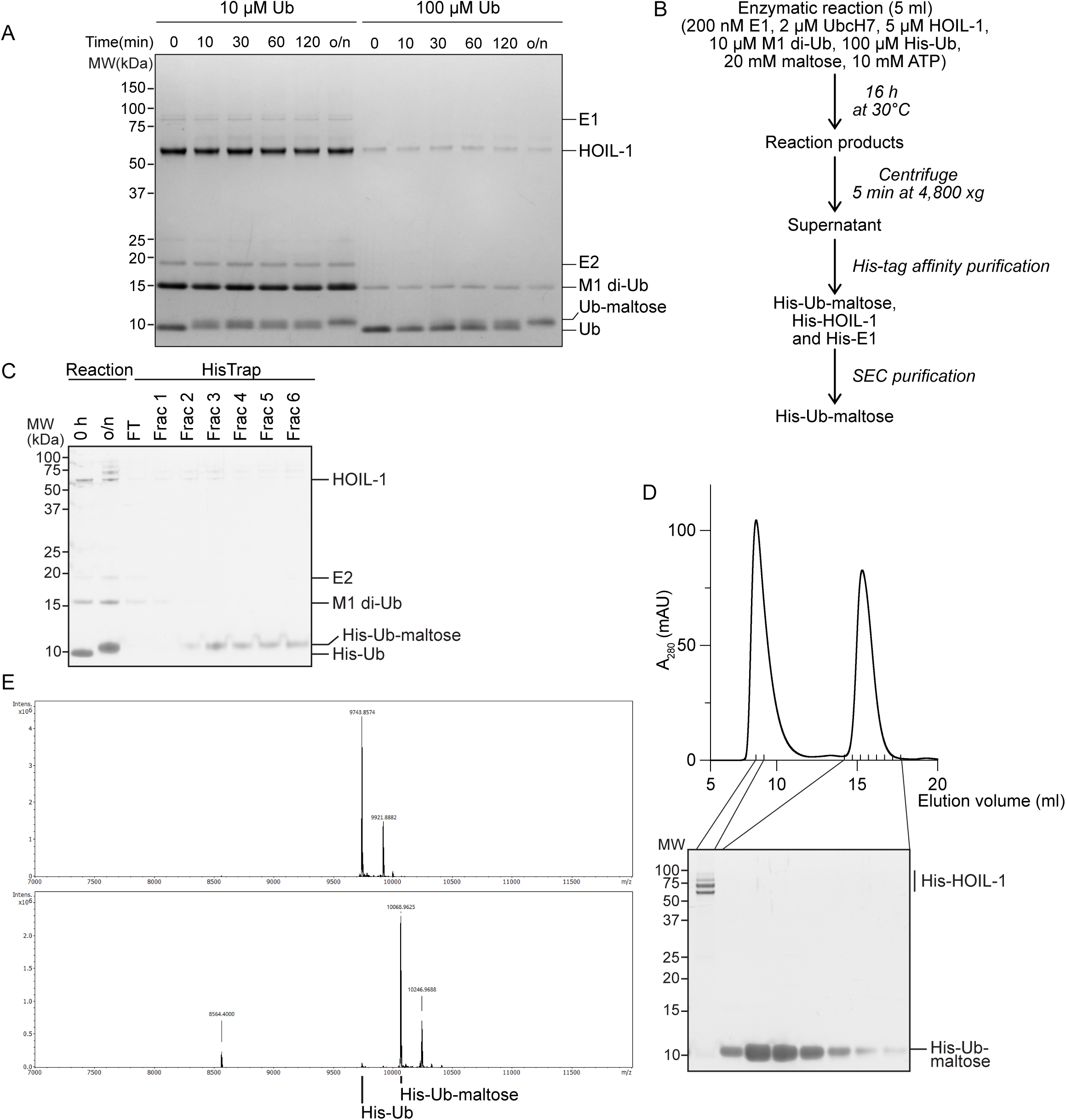
Large scale preparation and purification of ubiquitinated maltose. **(A)** Time-course showing HOIL-1-mediated ubiquitination of maltose using low (10 µM) and higher (100 µM) ubiquitin concentrations. To enable comparison of ubiquitin (Ub) and ubiquitin-maltose bands between the different samples, the samples containing 100 µM were diluted 10-fold before SDS-PAGE analysis. o/n, overnight. **(B)** Workflow for large scale ubiquitin-maltose preparation and purification. **(C)** Coomassie-stained SDS-PAGE gel showing successful large-scale ubiquitin-maltose generation (Reaction) and His-tag affinity purification step (HisTrap). **(D)** Size-exclusion chromatography purification of His-ubiquitin-maltose after HisTrap purification. (**E**) Intact mass spectrometry of His-ubiquitin (top) and purified His-ubiquitin-maltose (bottom). The mass difference of 325 Da confirms mono-ubiquitination of a single maltose molecule. The slightly heavier minor species in the two spectra likely represent α-*N*-6-phosphogluconoylation of the His-tag, often identified in His-tagged proteins expressed in *E. coli* (65).

### Application of ubiquitin-maltose to investigate non-proteinaceous ubiquitination *in vitro*

To demonstrate the utility of our ubiquitinated maltose tool biomolecule, we followed two strategies. First, we attempted to crystallise the HOIL-1 (C460A) RING2 domain with His-ubiquitin maltose to obtain insights into how HOIL-1 may ubiquitinate maltose. We used the same crystallisation conditions as described for HOIL-1 RING2/ubiquitin above and successfully determined the X-ray structure to 1.7 Å resolution (Table 1). As with the HOIL-1 RING2/ubiquitin structure, the crystallographic asymmetric unit contains two HOIL-1 RING2 molecules and one ubiquitin molecule bound in the donor ubiquitin site of one HOIL-1 RING2 domain (Fig EV4A, B). While we can resolve the ubiquitin C-terminus, we do not observe clear electron density for the ubiquitin-conjugated maltose molecule and therefore were unable to build the maltose molecule into the model (Fig EV4C). This suggests that maltose does not form specific stable interactions with the HOIL-1 RING2 domain, in line with the broad substrate specificity we observe for HOIL-1 *in vitro*.

In a separate approach, we used ubiquitin-maltose to investigate a small panel of DUBs to identify which DUBs can cleave the non-proteinaceous substrate ubiquitin-maltose. Our array consists of USP2, OTUD1, OTULIN, Cezanne and ATXN3. USP2 features broad substrate specificity that can cleave all seven Lys-linked ubiquitin chains (K6, K11, K27, K29, K33, K48 and K63), M1-linked ubiquitin and oxyester-linked Thr-ubiquitin (46, 47). The ubiquitin chain type-specific DUBs OTUD1 (K63-specific), OTULIN (M1-specific) and Cezanne (K11-specific) do not cleave ubiquitinated threonine, whereas ATXN3 does not cleave ubiquitinated lysine but can cleave ubiquitinated threonine (46, 47). As a positive control for oxyester-linked ubiquitin, we used hydroxylamine-treatment. We first verified DUB activity against a panel of amine-conjugated ubiquitin chains (K63-, K11- and M1-linked) and observed the expected activities for each DUB (Fig 5A, Fig EV5). When we incubated the same DUBs with the purified His-ubiquitin-maltose substrate, we observe that the non-specific USP2 can cleave ubiquitin-maltose, as can the positive control hydroxylamine, as expected (Fig 5A,B). Out of the chain-linkage specific DUBs, OTULIN is unable to cleave ubiquitin-maltose as expected (Fig 5A,B). Surprisingly, OTUD1 and Cezanne, which are not active against the Ub-threonine oxyester (47), can cleave the ubiquitin-maltose oxyester (Fig 5A,B). Finally, ATXN3 also can cleave the ubiquitin-maltose oxyester linkage (Fig 5A,B). These proof-of-principle experiments exemplify that our HOIL-1-catalysed ubiquitin-maltose can be utilised for enzymatic DUB assays *in vitro*. The observation that not all DUBs tested in our assay hydrolyse ubiquitin-maltose highlights that ubiquitin-maltose cleavage is a specific reaction. In line with this, OTUD1 and Cezanne, with reported poor activity against the oxyester-linked substrate ubiquitin-Thr (47), cleave ubiquitin-maltose in our assay.

**Figure 5:**
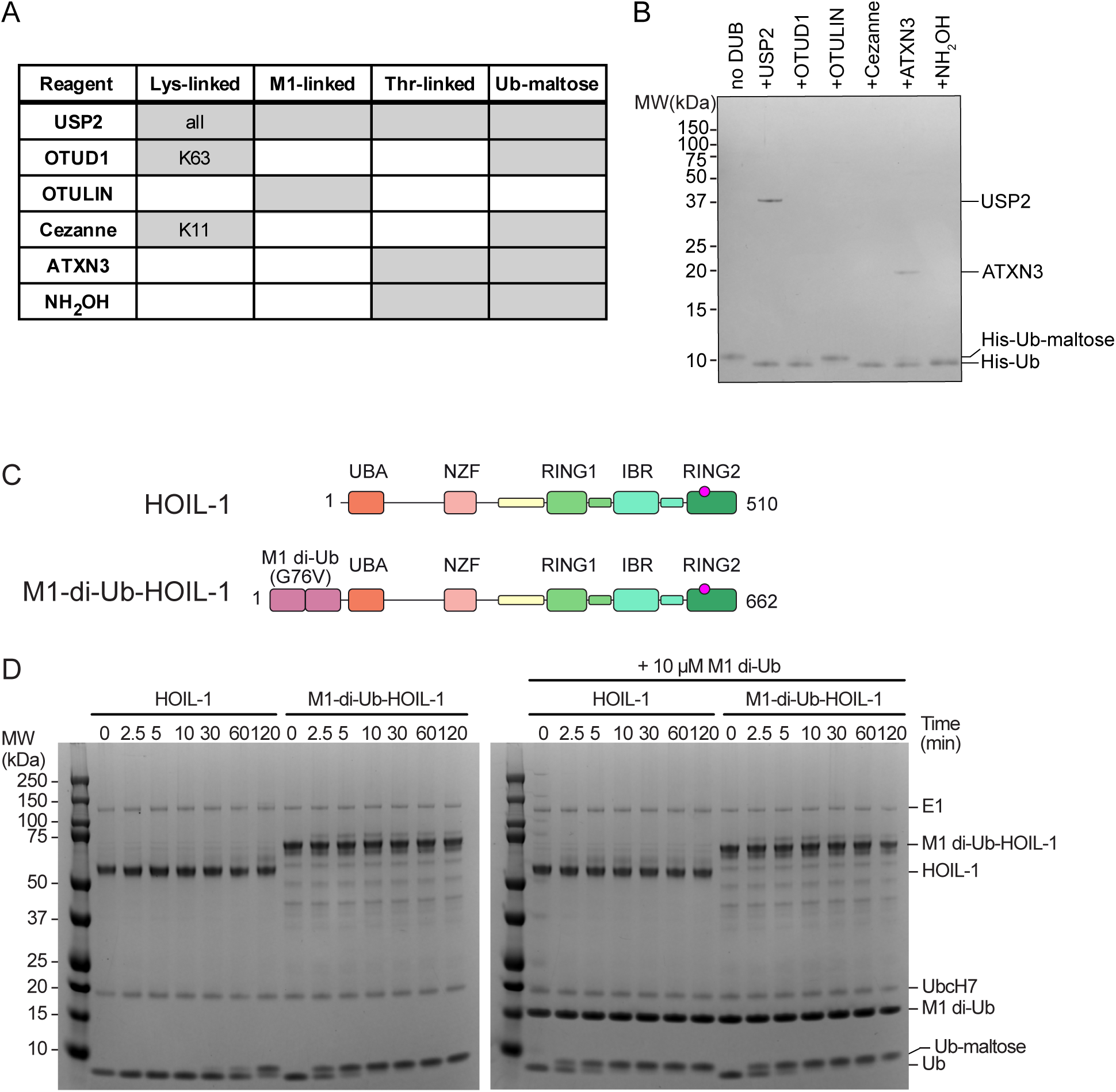
Proof-of-principle DUB assay towards His-ubiquitin-maltose hydrolysis and design of a constitutively active HOIL-1 variant. **(A)** Table summarising known activities of DUBs and hydroxylamine (NH_2_OH) against different amino acid ubiquitin linkages (Lys, M1, Thr) from the literature and results from our analysis with Ub-maltose. Grey cells indicate activity with the indicated species. For Lys chain-type specific DUBs, the cleaved linkages are indicated. **(B)** Ubiquitin-maltose DUB assay. His-ubiquitin-maltose was incubated with different DUBs as indicated and analysed via SDS-PAGE. The first lane contains untreated sample, the sample in the last lane was treated with hydroxylamine (NH_2_OH) to fully cleave oxyester-linked His-ubiquitin-maltose. DUBs were used at different, optimal concentrations (46), hence only DUBs used at high concentrations (USP2, ATXN3) are resolved on the gel. **(C)** Domain organisation of endogenous human HOIL-1 (top) and the constitutive active M1-di-Ub-HOIL-1 fusion protein that contains an N-terminal M1-linked di-ubiquitin in which glycine 76 in both protomers is mutated to valine (G76V). **(D)** Maltose-ubiquitination assay comparing HOIL-1 with the M1-di-Ub(G76V)-HOIL-1 fusion protein in the presence and absence of additional allosteric M1-linked di-ubiquitin. Ub: ubiquitin.

### Generation of a constitutively active HOIL-1 variant enables efficient maltose ubiquitination in the absence of allosteric M1-linked di-ubiquitin

HOIL-1 is fully active only in the presence of allosteric M1-linked di-ubiquitin (25, 29, 30), necessitating the addition of high concentrations of di-ubiquitin in our assay, or increasing the reaction times. This complicates our *in vitro* ubiquitination workflow, may introduce artifacts or may require additional purification steps to remove di-ubiquitin. Therefore, we aimed to generate a constitutively active HOIL-1 by fusing M1-linked di-ubiquitin to the HOIL-1 N-terminus (M1-di-Ub-HOIL-1, Fig 5C). A similar approach has previously been implemented in a cellular context for the RBR E3 ligase HOIP, which is also activated by M1 di-ubiquitin (48). We initially used the native human M1 di-ubiquitin protein sequence with different linker lengths between di-ubiquitin and HOIL-1. While we were able to recombinantly express these constructs in *E. coli*, we observed large amounts of cleaved M1 di-ubiquitin and free HOIL-1, which made it challenging to obtain a homogenous fraction of M1-di-Ub-HOIL-1 (Fig EV6A). In our final design, we therefore completely removed the linker between M1-di-ubiquitin and HOIL-1 and replaced Gly76 in both ubiquitin moieties with Val (G76V). This final construct expressed and purified almost as well as HOIL-1 (Fig EV6B). Importantly, when we compared the activity of the M1-di-Ub-HOIL-1 to HOIL-1 in the presence and absence of additional M1-linked di-ubiquitin, we observed that M1-di-Ub-HOIL-1 without exogenous M1-linked di-ubiquitin is as active as HOIL-1 in the presence of the allosteric activator and is not further activated by addition of exogenous M1-linked di-ubiquitin (Fig 5D). This experiment shows that M1-di-Ub-HOIL-1 is a constitutive active enzyme that can replace HOIL-1 and M1-linked di-ubiquitin to streamline our ubiquitin-maltose generation system.

## DISCUSSION

Ubiquitination of non-proteinaceous substrates including lipopolysaccharides, sugars, ADPR, ribonucleotides and lipids is a rapidly emerging theme in the ubiquitin field (21–26). The RBR E3 ligase HOIL-1 is one of the E3 ubiquitin ligases that has been implicated in ubiquitinating maltose and glycogen, in addition to its previously reported activity in ubiquitinating Ser, Thr and Lys residues in proteins (17–20, 25, 29, 30).

Our work here provides further quantitative insights on the catalytic mechanism of the RBR E3 ubiquitin ligase HOIL-1 and its relative substrate specificity. Using *in vitro* ubiquitination assays on model peptide substrates and an array of saccharides, we show that HOIL-1 has a strong preference for ubiquitinating hydroxyl groups in Ser sidechains, maltose and other saccharides with weaker activity for ubiquitinating Thr and no detectable activity for the Lys-containing model peptide.

Our biochemical analysis identifies the catalytic His510 residue in HOIL-1’s C-terminus as a critical factor for Ser ubiquitination. The HOIL-1 H510A mutant is unable to ubiquitinate the Ser model peptide, but, surprisingly, gains activity to ubiquitinate Lys residues in the model peptide and on HOIL-1 itself. Our crystal structure of the HOIL-1 RING2 domain with and without ubiquitin shows that His510 forms part a dynamic HOIL-1 catalytic centre that includes the loop containing the catalytic Cys460 and Trp462. Our structure demonstrates that ubiquitin-binding induces a rearrangement of these regions into what is likely the active conformation that is also observed in previous structures of the HOIL-1/E2-ubiquitin thioester transfer complex. Our biochemical and structural analyses, together with our previous observation (29) that His510 is dispensable for HOIL-1’s first catalytic step (ubiquitin transfer from E2 to E3), but critical for ubiquitin transfer from the E3 to the substrate, suggest that His510 is required to deprotonate and activate the incoming acceptor nucleophile. Compared to amines, hydroxyl groups are weaker nucleophiles and therefore critically require activation by HOIL-1’s His510 for efficient ubiquitin transfer to the substrate. Mutation of His510 to Ala removes this activating effect, thereby preventing ubiquitin transfer from the E3 active site to the substrate hydroxyl group. The E3-ubiquitin thioester then has two options: either discharge ubiquitin onto a solvent molecule (water/hydroxide ion) or to an amine, either on a Lys sidechain in HOIL-1 itself or a Lys in the model substrate (if present). This results in the apparent specificity switch from Ser to Lys for the H510A mutant compared to WT HOIL-1.

Our work further shows that HOIL-1 has broad substrate specificity against different mono- and di-saccharides. Previous publications show that HOIL-1 ubiquitinates the C6 hydroxymethyl (CH_2_OH) moiety in a glucose subunit of maltoheptaose (49). All successful substrates tested in our work contain a hydroxymethyl moiety, either in the Ser side chain or the various saccharides. In the saccharides, the CH_2_OH moiety is mostly found in the C6 position, with the exceptions of Kdo, where the hydroxymethyl moiety includes C8, and Neu5AC, where it includes C9 (Fig 3B,E). The major caveat of the *in vitro* experiments here and elsewhere (25, 29, 30) is that very high substrate concentrations of 20 mM are used, which may have limited physiological relevance. As we have argued before, just because an E3 ligase can ubiquitinate a substrate *in vitro*, does not mean that it does so in a cellular environment (41). On the flip side, high local concentrations of maltose subunits, as found in the proposed HOIL-1 substrate glycogen, may provide the right environment for efficient HOIL-1-mediated ubiquitination of these biomolecules in the cell. We also observe HOIL-1-mediated *in vitro* ubiquitination of saccharides found in high concentration on the surface of pathogens, e.g. Kdo, a component of bacterial LPS, and MurNAc, a component of bacterial peptidoglycan. This observation may hint at additional functions of HOIL-1 in ubiquitinating intracellular pathogens, as previously described for the E3 ligase RNF213 (21), and in line with LUBAC’s role in innate immune signalling. In all cases, to overcome the limitations of these *in vitro* experiments, it will be critical to develop novel technologies that can detect oxyester linked non-proteinaceous substrates, such as ubiquitinated sugars in the cell.

Recent publications report that current mass spectrometry methods may not be able to detect oxyester-conjugated ubiquitin species due to their reduced stability compared to isopeptide linked ubiquitinated species (15). Our approach to generate ubiquitinated model products will provide benchmarks that can be utilised to develop and optimise mass spectrometry workflows that are compatible with ubiquitinated sugars, e.g. by spiking-in our model products in cell lysates and detecting the expected masses. In addition, ubiquitinated sugars generated with our protocol can be used to investigate DUB activity and specificity as shown in our proof-of-principle work. Our experiments show that ubiquitin-maltose cleavage is a specific enzyme-catalysed reaction, as not all DUBs tested cleave the substrate. However, surprisingly, DUBs that previously were shown to be inactive against Ub-Thr oxyester bonds (OTUD1 and Cezanne, (47)), can cleave the ubiquitin-maltose oxyester. This highlights the specificity of the oxyester hydrolysis and substrate-specificity of these DUBs and the importance of comparing various oxyester-containing species as we have performed here for HOIL-1. To further delineate the differences in DUB activities against various oxyester-linked ubiquitin species, more detailed experiments with different enzyme concentrations and time-courses are required in the future. Nevertheless, our proof-of-principle study provides a strong foundation for these experiments. In addition to the assays presented here, we envision that our HOIL-1-generated ubiquitin-saccharide species will be valuable for investigating other aspects of non-proteinaceous ubiquitination, for example to pull down specific ubiquitin-sugar binding proteins which could then be identified by proteomics methods, supplementing other emerging technologies specific for oxyester-linked ubiquitin (50).

In summary, our work provides novel insights on the catalytic activity of the RBR E3 ubiquitin ligase HOIL-1 and its non-canonical ubiquitination activity towards non-proteinaceous substrates. We highlight how HOIL-1 and an engineered constitutively active HOIL-1 variant can be used to generate tool compounds and standards to interrogate non-proteinaceous ubiquitination and to develop urgently needed techniques in this burgeoning field.

## METHODS

### Plasmids and reagents

The plasmid for bacterial expression of full-length human His-3C-HOIL-1 (pOPINB-HOIL-1 full-length) is available from Addgene (Plasmid #193858) (29). His-3C-HOIL-1 RING2 domain (residues 412 – 510) in pOPINB was cloned from this plasmid. UBE1, UbcH7, native ubiquitin, non-cleavable His-ubiquitin and M1 di-ubiquitin constructs were described previously (29, 51, 52). The M1-di-Ub(G76V)-HOIL-1 fusion construct was cloned via in-fusion cloning by introducing the sequence of M1-di-Ub(G76V) into the pOPINB-HOIL-1 full-length plasmid. The pOPINB-M1-di-Ub(G76V)-HOIL-1 fusion plasmid for bacterial expression is available from Addgene (Plasmid # 229539). Saccharides were purchased from Biosynth Carbosynth (UK). Tn, sTn and T glycosylamino acids were synthesised according to a method described by Corcilius *et al*. (53).

### Protein expression and purification

Full-length HOIL-1 was expressed in *E. coli* Rosetta(DE3)pLacI, and all other recombinant proteins including the M1-di-Ub(G76V)-HOIL-1 fusion were expressed in *E. coli* BL21(DE3). Bacterial cultures were grown in 2x YT media with antibiotics until OD_600_ ∼0.8. Protein expression was induced with 0.5 mM IPTG (Golden Biotechnology). Cultures expressing HOIL-1 protein were additionally supplemented with 0.5 mM ZnCl_2_ at time of induction. The media for the M1-di-Ub(G76V)-HOIL-1 fusion expression was additionally supplemented with 15 mL of 750 mM mannitol per litre of media to reduce lower molecular weight contaminants found during purification. Cultures were grown overnight at 20°C before harvesting by centrifugation.

For purification, cells were lysed with addition of lysozyme and sonication in the presence of protease inhibitors (PMSF and leupeptin), DNaseI and MgCl_2_. Cell lysates were cleared by centrifugation. To purify His-tagged E2, ubiquitin, M1 di-ubiquitin and full-length and RING2 HOIL-1 proteins, cleared cell lysates were applied to Ni-NTA His-bind resin (Merk Millipore) equilibrated in gravity columns. His-tagged proteins were eluted with high salt purification buffer (50 mM Tris pH8, 500 mM NaCl, 10% sucrose and 10% glucose) supplemented with 300 mM imidazole. Eluted proteins were dialysed overnight in the presence of 1/100 (v/v) His-HRV 3C protease to remove the His-tag. Cleaved proteins were applied to Ni-NTA His-bind resin to remove His-HRV 3C protease before purification by size-exclusion chromatography (Superdex 75 16/600 pg or Superdex 200 16/600 pg, Cytiva) using the AKTA pure chromatography system (Cytiva).

The His-3C-M1-di-Ub(G76V)-HOIL-1 fusion protein, was purified basically as described above for full-length HOIL-1 but with an additional wash step with 30 mM imidazole in high salt purification buffer before elution in high salt purification buffer supplemented with 300 mM imidazole and the His-tag was not cleaved.

K11 tri-ubiquitin and K63 tetra-ubiquitin were generated enzymatically from mono ubiquitin with ubiquitin linkage-specific enzymes as described (54). The DUBs USP2, OTUD1, OTULIN, Cezanne and ATXN3 were expressed in *E. coli* and purified as previously described (46).

All purified proteins were concentrated and snap frozen in liquid nitrogen before storing at - 80°C.

### Peptide ubiquitination assays

A reaction master mix was made with 200 nM E1, 2 μM UbcH7, 5 μM HOIL-1, 10 μM blocked M1 di-ubiqutin and 10 μM native ubiquitin in DPBS buffer (Gibco) containing 10 mM MgCl_2_ and 0.5 mM TCEP. 200 mM peptide stocks (GenScript Biotech, Singapore, 20 mM final concentration) were mixed with 0.1 M ATP (pH 7, 10 mM final concentration) and added to the master mix to initiate the reaction. Reactions were incubated at 30°C and sampled at indicated time points by mixing with 1x LDS sample buffer (NuPAGE) supplemented with 0.1 M DTT. Reactions in Figure 2 were further treated with 1.5 M NH_2_OH (Sigma-Aldrich 467804) as indicated for 30 min at 30°C before addition of 1x LDS sample buffer supplemented with 0.1 M DTT. All samples were denatured at room temperature for 5 min and analysed by SDS-PAGE followed by Coomassie staining. Gels were imaged on a ChemiDoc MP Imaging system (Bio-Rad) and band intensities quantified with Image Lab (Bio-Rad). Plots were generated in Prism 10 (Graphpad).

### Saccharide ubiquitination assays

A reaction master mix was made with 200 nM E1, 2 μM UbcH7, 5 μM HOIL-1, 10 μM blocked M1 di-ubiquitin and 10 μM native ubiquitin in DPBS buffer (Gibco) containing 10 mM MgCl_2_ and 0.5 mM TCEP. 200 mM stocks of individual saccharides dissolved in DPBS buffer were added into individual reactions to reach 20 mM final concentration. 0 min samples were taken by mixing with 1x LDS sample buffer (NuPAGE) supplemented with 0.1 M DTT. 10 mM ATP (pH 7) was added to initiate the reactions. Reaction samples were incubated at 30°C while indicated time point samples were taken by mixing with 1x LDS buffer supplemented with 0.1 M DTT. All samples were denatured at room temperature for 5 min and analysed by SDS-PAGE followed by Coomassie staining.

The assays for the physiological saccharides were performed in a similar manner, but without the addition of the allosteric activator M1 di-ubiquitin and with increased reaction times as indicated. Samples for MALDI-TOF analysis were taken at 0, 2 and 4 h by acidification with 0.4% TFA (Sigma Aldrich).

### Mass spectrometry

#### MALDI-TOF

A previously described protocol was followed to prepare MALDI samples (55). In brief, acidified sample was diluted 1:1 v/v with ACN/FA solution (acetonitrile mixed with a 5% v/v formic acid in water in a 70:30 v/v ratio). The MALDI target was pre-treated with a thin layer of saturated α-CHCA (*α*-cyano-4-hydroxycinnamic acid) dissolved in ACN/FA solution. 20 mg/mL DHB solution was prepared in ACN/TFA solution (acetonitrile mixed with 0.1% v/v trifluoroacetic acid in water in a 70:30 v/v ratio). 0.5 μl diluted sample were deposited onto the target followed by 0.5 μL α-CHCA/DHB solution (1:1 v/v) and air dried. Data were acquired on a Bruker microflex using 100% laser power with no detection gain. Data were automatically analysed by the software flexAnalysis 3.4 (Bruker) and replotted using Prism 9 (Graphpad).

#### Intact MS

Purified His-ubiquitin and His-ubiquitin-maltose were diluted to 20 μΜ using milliQ H_2_O. Peptide ubiquitination samples were acidified with sodium acetate solution (pH 4.6) to terminate reactions. All samples were centrifugated for 5 min at 13,000 x g and 4°C to remove aggregates. 5 μl of His-ubiquitin and His-ubiquitin-maltose and 15 μl of peptide ubiquitination samples were directly infused onto a Maxis II (Bruker) mass spectrometer at 200°C with 4 l/min drying gas and a capillary voltage of 2,000 V. Data were acquired for 1 min at 300-3000 m/z, and the data was deconvoluted using the maximum entropy algorithm within the Bruker Data Analysis software.

### DUB assay

Indicated DUBs were pre-diluted and incubated with DUB dilution buffer containing 25 mM Tris pH7.5, 150 mM NaCl and 10 mM DTT. M1 di-ubiquitin, K11 tri-ubiquitin and K63 tetra-ubiquitin were mixed with DUB reaction buffer containing 50 mM Tris pH7.5, 50 mM NaCl and 5 mM DTT. Similarly, His-ubiquitin-maltose was mixed with DUB reaction buffer. To set up the control reaction, the polyubiquitin mixture containing 5 μΜ of each chain type was split and mixed with individual DUBs. The final concentration for each DUB is USP2 2 μΜ, OTUD1 0.2 μΜ, OTULIN 0.1 μΜ, Cezanne 0.2 μΜ and ATXN3 2 μΜ. DUBs were mixed at the same concentration with 5 μΜ His-ubiquitin-maltose. All reactions were incubated at RT for 30 min. Reactions were stopped by adding LDS buffer supplemented with DTT. Samples were resolved by SDS-PAGE followed by Coomassie staining.

### Large scale ubiquitinated saccharide production and purification

To enzymatically prepare ubiquitin-maltose, a reaction containing 200 nM E1, 2 μΜ UbcH7, 5 μΜ ΗΟIL-1, 10 μM blocked M1 di-ubiquitin, 100 μM His-ubiquitin and 20 mM maltose was made up in DPBS buffer supplemented with 10 mM MgCl_2_ and 0.5 mM TCEP in 5 ml final volume. The reaction was initiated by addition of 10 mM ATP, incubated at 30°C and sampled as indicated to check on reaction completeness by SDS-PAGE. After overnight (∼16 h) incubation, the reaction solution was clarified by centrifugation (5 min at 4,800 x g) and supernatant was applied to a 1 ml His-Trap HP column (Cytiva) equilibrated in DPBS. Bound resin was washed with DPBS buffer and block-eluted with DPBS buffer containing 300 mM imidazole. The eluates were examined by SDS-PAGE, pooled and concentrated for further size-exclusion chromatography purification on a Superdex 75 Increase 10/300 column (Cytiva) equilibrated in DPBS. Purified His-ubiquitin-maltose was concentrated, aliquoted and snap frozen in liquid nitrogen then stored at -80 °C.

### Crystallisation

Purified HOIL-1 helix-RING2 (residues 412-510 with the C460A mutation) was mixed with His-tagged ubiquitin or His-tagged ubiquitin-maltose in a 1:2 molar ratio. The protein mixture was concentrated using Amicon Ultra 3 kDa MW cut-off device (Merck Millipore) to ∼15 mg/ml and used to set up commercial crystallisation screens using the sitting drop vapour diffusion method at room temperature. Initial crystal hits were obtained from the Natrix (Hampton) screen in a condition containing 0.01 M MgCl_2_, 0.05 M MES pH5.6 and 1.8 M LiSO_4_ by mixing 100 nl protein with 100 nl reservoir solution. Hit crystals were optimised by titrating MgCl_2_ (0.01 – 0.04 M) as well as varying MES pH from 5.2 to 6.2. The best diffracting crystals grew in a pH range from 5.2 – 6.0 and various MgCl_2_ concentrations. The HOIL-1 helix-RING2/Ub-maltose structure was determined from a crystal grown in 0.02 M MgCl_2_, 0.05 M HEPES pH6 and 1.8 M LiSO_4_. The crystal was dehydrated by replacing the reservoir solution with 2.5 M LiSO_4_ and incubation overnight. The HOIL-1 helix-RING2/Ub structure was determined from a crystal grown in 0.01 M MgCl_2_, 0.05 M HEPES pH6.2 and 1.8 M LiSO_4_ All crystals were rinsed in fresh reservoir solution and directly flash frozen in liquid nitrogen.

### Diffraction data collection, processing, model building and refinement

Crystal diffraction data were collected at the Australian Synchrotron MX2 beamline (56) at a wavelength of 0.9537 Å and 100 K temperature. Datasets were indexed and integrated with XDS (57) and merged with AIMLESS (58) using the automated processing pipeline at the Australian Synchrotron. Phaser (59) was used to solve the structure by molecular replacement. The initial search was performed using two copies of HOIL-1 RING2 (residues 444–510) molecules from the known HOIL-1 structure (PDB: 8EAZ) (29) and ubiquitin (PDB: 1UBQ) (60) each. Two HOIL-1 and one ubiquitin molecules were successfully found. The RING2 N-terminal helix of chain A (residue 425-443) and chain B (residue 431-443) were manually built. Iterative cycles of model-building and refinement were completed using Coot (61) and Phenix.refine (62, 63). Date refinement and statistics are shown in Table 1. All structural figures were prepared using UCSF ChimeraX (64).

## DATA AVAILABILITY

Atomic structures and diffraction data have been deposited in the PDB with accession codes 9EGV (HOIL-1 RING2/Ub) and 9EGW (HOIL-1 RING2/Ub-maltose). All other data supporting the conclusions are available in the article.

## ACKNOWLEDGEMENTS

We thank other members of the Lechtenberg and Komander labs for helpful discussions and suggestions and Geoffrey Kong (Monash Macromolecular Crystallisation Facility) for help with crystallisation. This work was funded by WEHI and National Health and Medical Research Council (NHMRC) Ideas Grants 1182757 (to B.C.L) and 2027601 (to E.D.G.-B.); Investigator Grants 1174941 (to R.J.P.), 2033340 (to E.D.G.-B.), 1178122 (to D.K.) and 2016268 (to B.C.L.), a Rebecca Cooper Fellowship (to E.D.G.-B.) and Australian Research Council DECRA Fellowship DE230100634 (to L.C.). X.S.W. is supported by an Australian Government Research Training Program Scholarship. The mass spectrometry analysis was performed at the WEHI Proteomics Facility. This research was undertaken in part using the MX2 beamline at the Australian Synchrotron, part of the Australian Nuclear Science and Technology Organisation (ANSTO), and made use of the Australian Cancer Research Foundation (ACRF) detector.

## AUTHOR CONTRIBUTIONS

X.S.W., A.C., M.J., J.J. and S.A.C. designed and performed experiments and interpreted data. E.D.G.-B. provided reagents and expertise on systematically exploring di-saccharide ubiquitination. K.H.T.M., L.C. and R.J.P. synthesised glycan reagents. D.K. provided reagents and ideas. B.C.L. designed the overall study, supervised the work, interpreted the results, secured funding and wrote the manuscript with input from all authors.

## MATERIALS & CORRESPONDENCE

Correspondence and requests for materials should be addressed to Bernhard C. Lechtenberg.

---

**Expanded View Figure 1:**
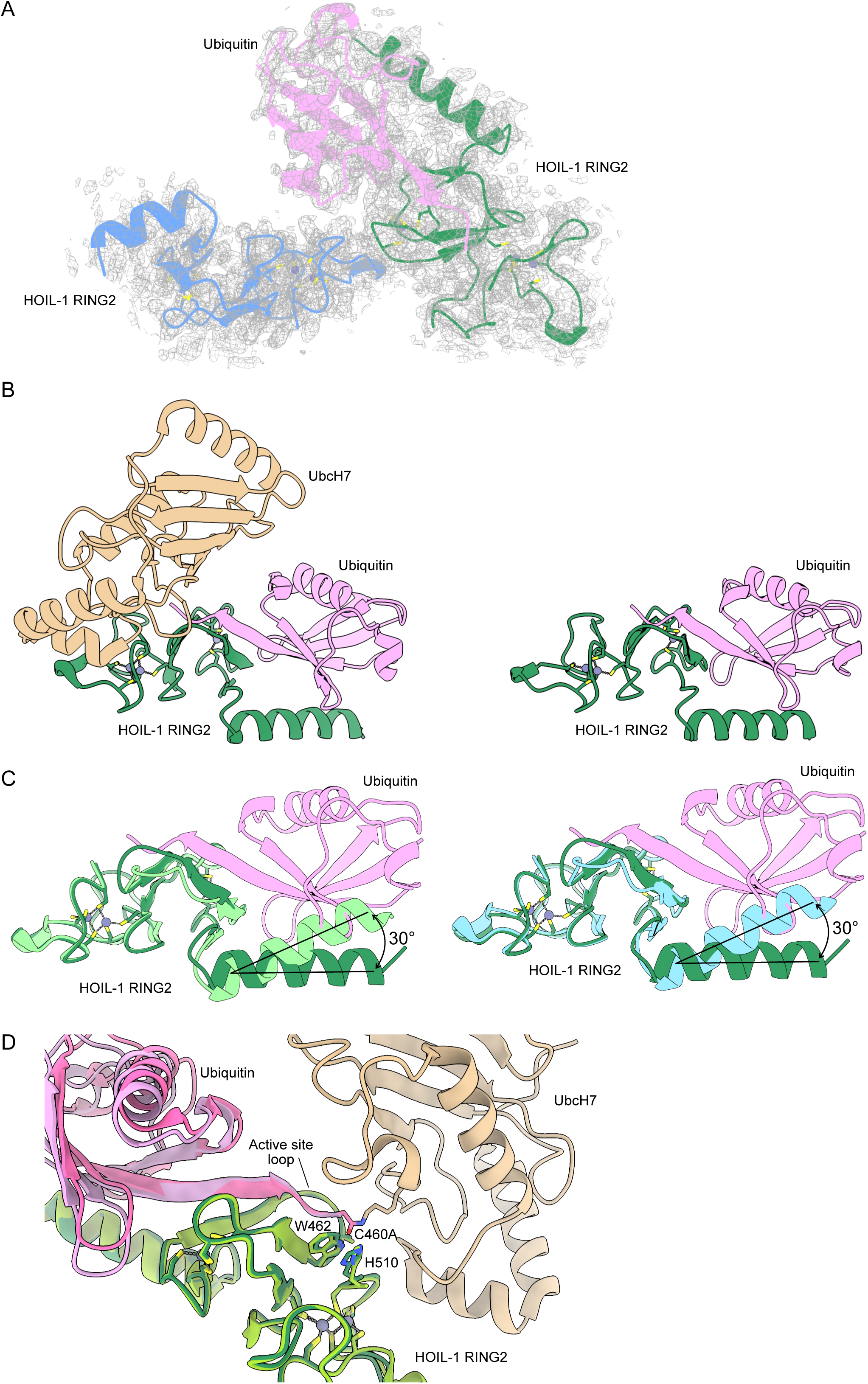
Crystal structure of HOIL-1 RING2/ubiquitin and comparison with other HOIL-1 RING2 structures. **(A)** Asymmetric unit of the HOIL-1 RING2 (green, blue) structure with ubiquitin (pink) with the 2Fo-Fc electron density (grey mesh) shown contoured at 1s. **(B)** Comparison of the UbcH7-ubiquitin conjugate-bound HOIL-1 RING2 domain (left, UbcH7 in wheat, HOIL-1 RING2 in green, ubiquitin in pink, PDB ID: 8EAZ, ref.(29)) and the HOIL-1 RING2/ubiquitin complex described here. **(C)** Overlays of the HOIL-1 RING2/ubiquitin complex described here (HOIL-1 in dark green, ubiquitin in pink) with two different free HOIL-1 RING2 domain structures. Left: PDB ID 7YUJ (light green) (30), Right: PDB ID 8BVL (cyan) (28). (D) Overlay of the HOIL-1 RING2/ubiquitin complex (HOIL-1 in dark green, ubiquitin in pink) and E2-ubiquitin conjugate-bound HOIL-1 RING2 (PDB ID: 8EAZ, ref.(29), HOIP in light green, ubiquitin in bright pink, UbcH7 in wheat).

**Expanded View Figure 2:**
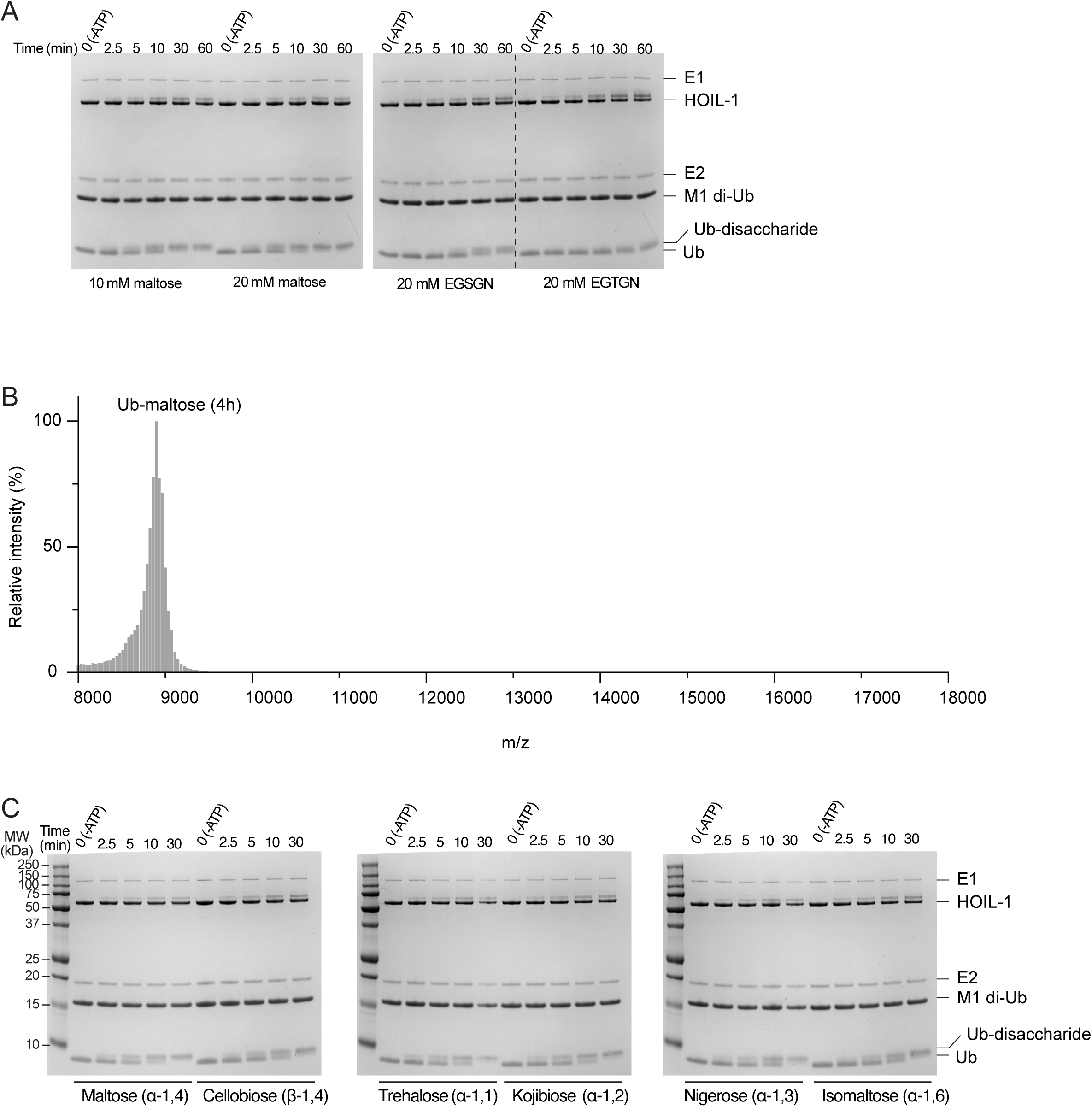
*In vitro* ubiquitination of peptides and di-saccharides by HOIL-1. **(A)** Coomassie-stained SDS-PAGE gels showing HOIL-1 ubiquitination of different substrates as indicated. Ubiquitin (Ub) and Ub-substrate bands were quantified for the graph in Figure 3A. Exemplary gels of three experiments are shown. **(B)** MALDI-TOF mass spectrometry of ubiquitinated maltose shows the expected mass of ubiquitin-maltose (8889 Da) with no evidence for dual modified maltose. **(C)** Coomassie-stained SDS-PAGE gels showing HOIL-1 ubiquitination of different di-saccharides as indicated. Ub-substrate and Ub bands were quantified for the graph in Figures 3C and 3D. Exemplary gels of three experiments are shown.

**Expanded View Figure 3:**
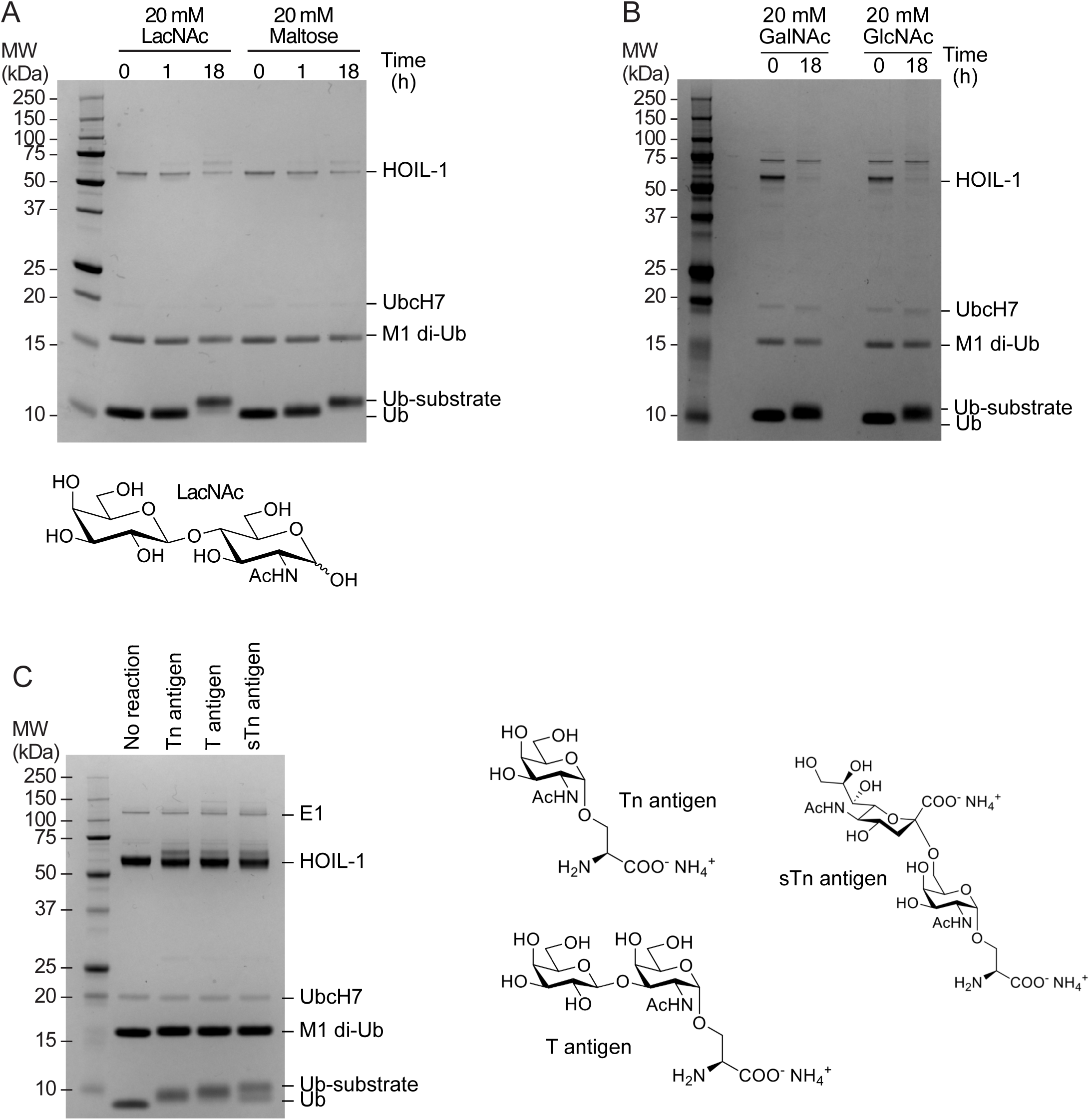
Examples of other ubiquitinated non-proteinaceous biomolecules generated by HOIL-1 *in vitro*. **(A)** Ubiquitination of N-acetyllactosamine (LacNAc) and maltose control. The chemical structure of LacNAc is shown. **(B)** Ubiquitination of GalNAc and GlcNAc. **(C)** Ubiquitination of glycosylated serine bearing the Tn antigen, T antigen and sTn antigen after 18 h. Chemical structures of the different substrates are shown on the right. Coomassie-stained SDS-PAGE gels are shown.

**Expanded View Figure 4:**
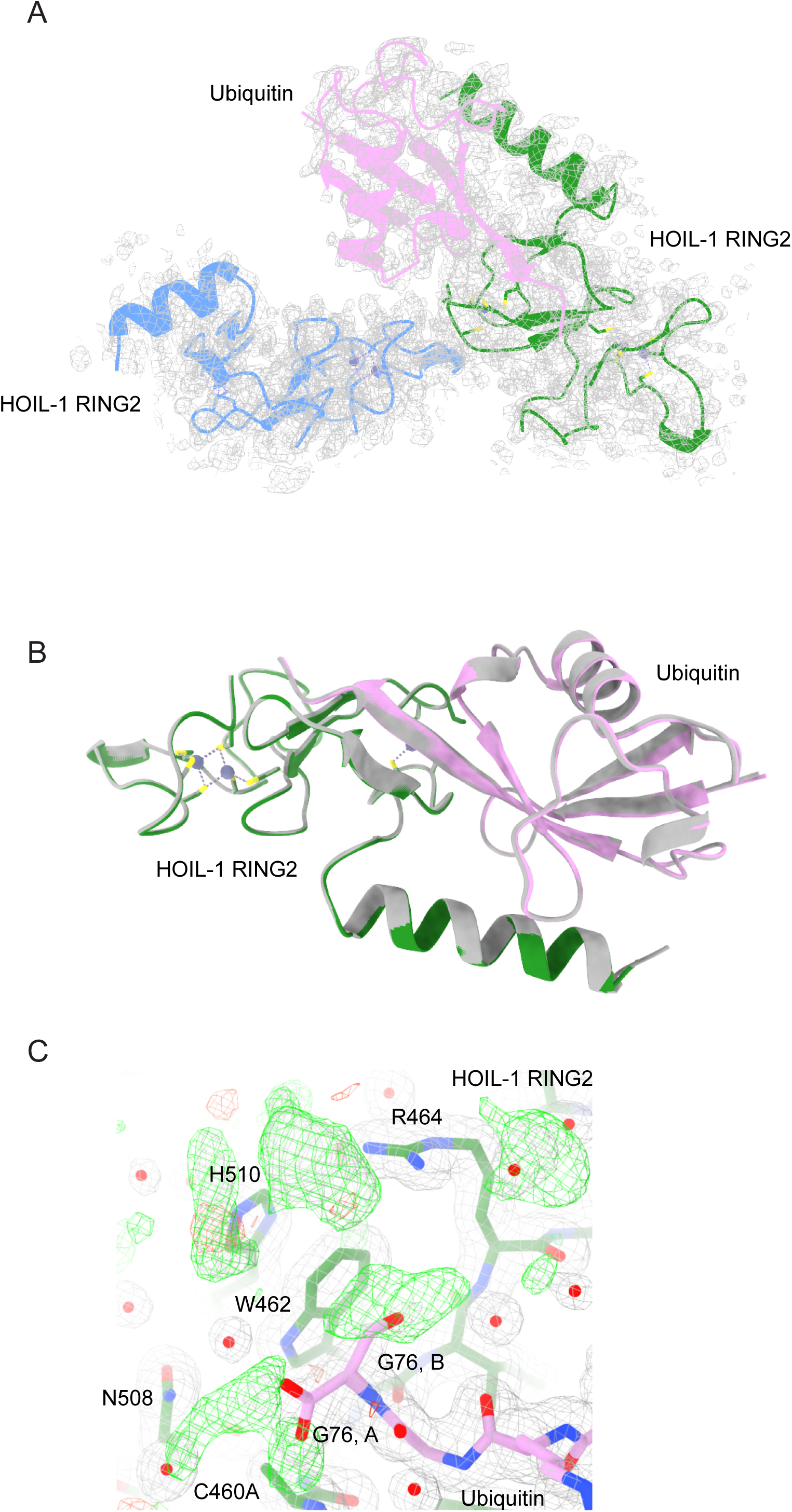
Crystal structure of HOIL-1 RING2/ubiquitin-maltose. **(A)** Asymmetric unit of the HOIL-1 RING2 (green, blue) structure with ubiquitin-maltose (pink) with the 2Fo-Fc electron density (grey mesh) shown contoured at 1σ. **(B**) Structural comparison of HOIL-1 RING2/ubiquitin-maltose (coloured as in A) and HOIL-1 RING2/ubiquitin (grey). **(C)** Electron density maps of the ubiquitin C-terminus (pink sticks; Gly76 modelled in two conformations, A and B) in the HOIL-1 (green sticks) active site lacks clear density for the maltose molecule. 2Fo-Fc map (grey mesh) shown contoured at 1σ and Fo-Fc difference density map (green and red mesh) shown contoured at +/- 2.5σ. Key residues are labelled. Water molecules are shown as red spheres.

**Expanded View Figure 5:**
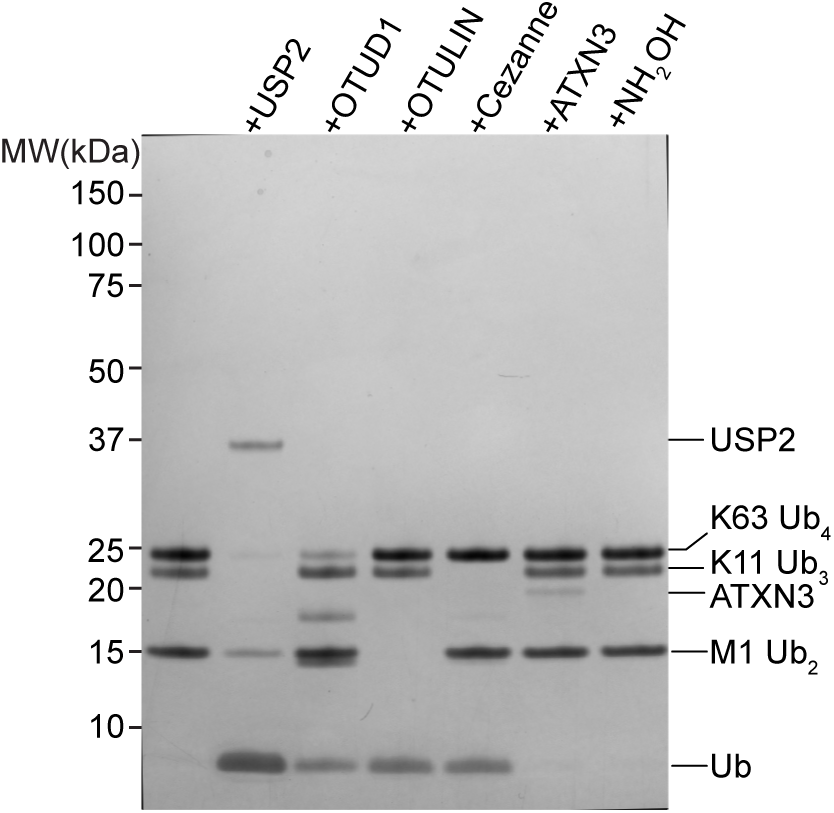
Quality control of the DUBs used for the ubiquitin-maltose DUB assay. To verify that DUBs used in Figure 5 are active and specific at the concentrations used, we tested their activity against a mixture of M1 di-ubiquitin, K11 tri-ubiquitin and K63 tetra-ubiquitin. As expected, USP2 cleaves all linkages, OTUD1 only K63 tetra-ubiquitin, OTULIN only M1 di-ubiquitin and Cezanne only K11 tri-ubiquitin. The DUB ATXN3 and the chemical hydroxylamine (NH_2_OH) do not cleave any of these (iso-)peptide linked ubiquitin oligomers. Ub: ubiquitin.

**Expanded View Figure 6:**
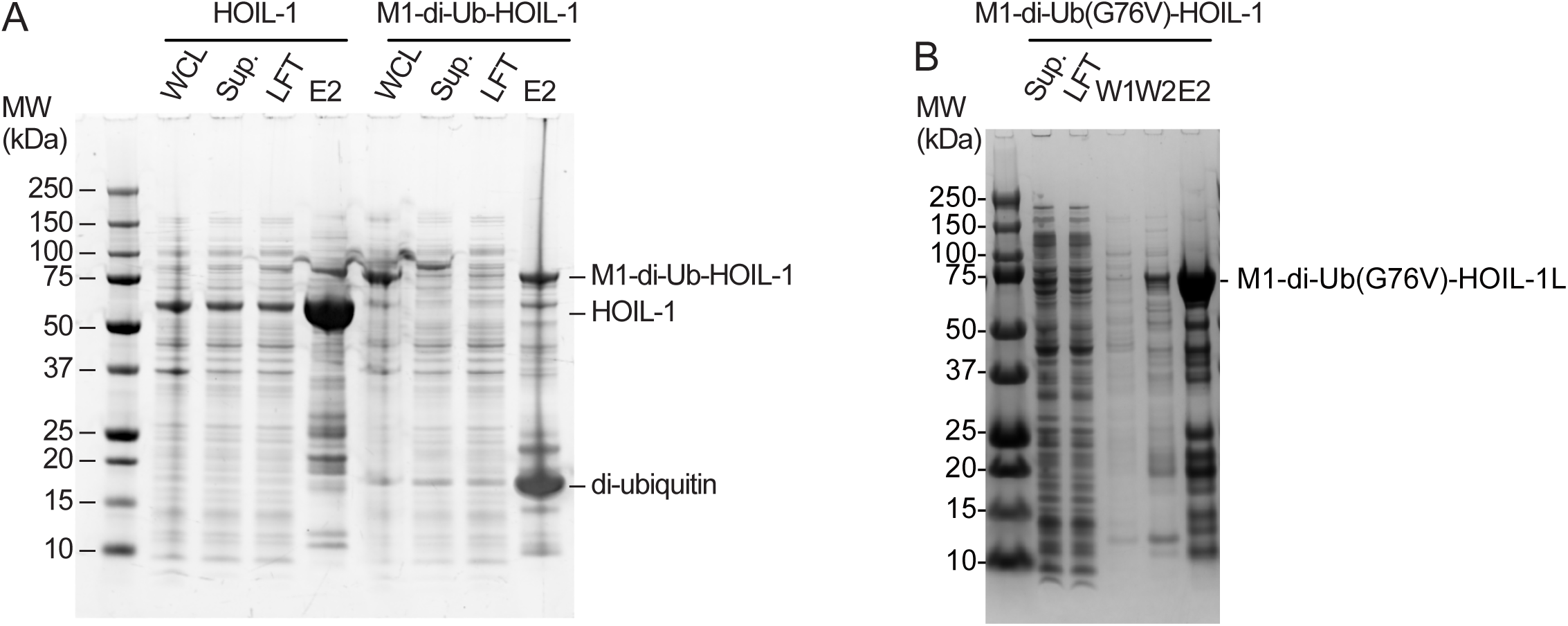
Design of a constitutively active HOIL-1 variant. **(B**) Initial purification (HisTrap) comparing WT human HOIL-1 and the first generation of the M1-di-Ub-HOIL-1 fusion protein. While both proteins express well in *E. coli* as indicated by the relevant bands in the whole-cell lysate (WCL), M1-di-Ub-HOIL-1 is not stable during HisTrap purification indicated by a large amount of di-ubiquitin in the HisTrap elution (E2) sample. **(C)** HisTrap purification of linker-less M1-di-Ub(G76V)-HOIL-1 showing that removal of the linker between the di-ubiquitin and HOIL-1 and mutation of Gly76 to Val (G76V) in both copies of ubiquitin yields a more stable fusion protein. For the assay shown in Fig 5D, the protein was further purified via size-exclusion chromatography (see Methods). Sup.: supernatant; LFT: load flow-through; W1: wash 1; W2: wash 2.

## REFERENCES

1. Hershko, A., and Ciechanover, A. (1998) The ubiquitin system Annu Rev Biochem 67, 425–479 10.1146/annurev.biochem.67.1.425

2. Komander, D., and Rape, M. (2012) The ubiquitin code Annu Rev Biochem 81, 203–229 10.1146/annurev-biochem-060310-170328

3. Kirisako, T., Kamei, K., Murata, S., Kato, M., Fukumoto, H., Kanie, M., Sano, S., Tokunaga, F., Tanaka, K., and Iwai, K. (2006) A ubiquitin ligase complex assembles linear polyubiquitin chains The EMBO journal 25, 4877–488710.1038/sj.emboj.7601360

4. Gerlach, B., Cordier, S. M., Schmukle, A. C., Emmerich, C. H., Rieser, E., Haas, T. L., Webb, A. I., Rickard, J. A., Anderton, H., Wong, W. W., Nachbur, U., Gangoda, L., Warnken, U., Purcell, A. W., Silke, J., and Walczak, H. (2011) Linear ubiquitination prevents inflammation and regulates immune signalling Nature 471, 591–596 10.1038/nature09816

5. Ikeda, F., Deribe, Y. L., Skanland, S. S., Stieglitz, B., Grabbe, C., Franz-Wachtel, M., van Wijk, S. J., Goswami, P., Nagy, V., Terzic, J., Tokunaga, F., Androulidaki, A., Nakagawa, T., Pasparakis, M., Iwai, K., Sundberg, J. P., Schaefer, L., Rittinger, K., Macek, B., and Dikic, I. (2011) SHARPIN forms a linear ubiquitin ligase complex regulating NF-kappaB activity and apoptosis Nature 471, 637–641 10.1038/nature09814

6. Tokunaga, F., Nakagawa, T., Nakahara, M., Saeki, Y., Taniguchi, M., Sakata, S., Tanaka, K., Nakano, H., and Iwai, K. (2011) SHARPIN is a component of the NF-kappaB-activating linear ubiquitin chain assembly complex Nature 471, 633–636 10.1038/nature09815

7. Smit, J. J., Monteferrario, D., Noordermeer, S. M., van Dijk, W. J., van der Reijden, B. A., and Sixma, T. K. (2012) The E3 ligase HOIP specifies linear ubiquitin chain assembly through its RING-IBR-RING domain and the unique LDD extension The EMBO journal 31, 3833–3844 10.1038/emboj.2012.217

8. Stieglitz, B., Morris-Davies, A. C., Koliopoulos, M. G., Christodoulou, E., and Rittinger, K. (2012) LUBAC synthesizes linear ubiquitin chains via a thioester intermediate EMBO Rep 13, 840–846 10.1038/embor.2012.105

9. Stieglitz, B., Rana, R. R., Koliopoulos, M. G., Morris-Davies, A. C., Schaeffer, V., Christodoulou, E., Howell, S., Brown, N. R., Dikic, I., and Rittinger, K. (2013) Structural basis for ligase-specific conjugation of linear ubiquitin chains by HOIP Nature 503, 422–426 10.1038/nature12638

10. Kelsall, I. R. (2022) Non-lysine ubiquitylation: Doing things differently Front Mol Biosci 9, 1008175 10.3389/fmolb.2022.1008175

11. Squair, D. R., and Virdee, S. (2022) A new dawn beyond lysine ubiquitination Nat Chem Biol 18, 802–811 10.1038/s41589-022-01088-2

12. Dikic, I., and Schulman, B. A. (2023) An expanded lexicon for the ubiquitin code Nat Rev Mol Cell Biol 24, 273–287 10.1038/s41580-022-00543-1

13. Pao, K. C., Wood, N. T., Knebel, A., Rafie, K., Stanley, M., Mabbitt, P. D., Sundaramoorthy, R., Hofmann, K., van Aalten, D. M. F., and Virdee, S. (2018) Activity-based E3 ligase profiling uncovers an E3 ligase with esterification activity Nature 556, 381–385 10.1038/s41586-018-0026-1

14. Purser, N., Tripathi-Giesgen, I., Li, J., Scott, D. C., Horn-Ghetko, D., Baek, K., Schulman, B. A., Alpi, A. F., and Kleiger, G. (2023) Catalysis of non-canonical protein ubiquitylation by the ARIH1 ubiquitin ligase The Biochemical journal 480, 1817–183110.1042/BCJ20230373

15. Yoshida, Y., Takahashi, T., Ishii, N., Matsuo, I., Takahashi, S., Inoue, H., Endo, A., Tsuchiya, H., Okada, M., Ando, C., Suzuki, T., Dohmae, N., Saeki, Y., Tanaka, K., and Suzuki, T. (2024) Sugar-mediated non-canonical ubiquitination impairs Nrf1/NFE2L1 activation Molecular cell 84, 3115–3127 e3111 10.1016/j.molcel.2024.07.013

16. Smit, J. J., van Dijk, W. J., El Atmioui, D., Merkx, R., Ovaa, H., and Sixma, T. K. (2013) Target specificity of the E3 ligase LUBAC for ubiquitin and NEMO relies on different minimal requirements The Journal of biological chemistry 288, 31728–31737 10.1074/jbc.M113.495846

17. Fuseya, Y., Fujita, H., Kim, M., Ohtake, F., Nishide, A., Sasaki, K., Saeki, Y., Tanaka, K., Takahashi, R., and Iwai, K. (2020) The HOIL-1L ligase modulates immune signalling and cell death via monoubiquitination of LUBAC Nat Cell Biol 22, 663–673 10.1038/s41556-020-0517-9

18. Rodriguez Carvajal, A., Grishkovskaya, I., Gomez Diaz, C., Vogel, A., Sonn-Segev, A., Kushwah, M. S., Schodl, K., Deszcz, L., Orban-Nemeth, Z., Sakamoto, S., Mechtler, K., Kukura, P., Clausen, T., Haselbach, D., and Ikeda, F. (2021) The linear ubiquitin chain assembly complex (LUBAC) generates heterotypic ubiquitin chains Elife 10, 10.7554/eLife.60660

19. McCrory, E. H., Akimov, V., Cohen, P., and Blagoev, B. (2022) Identification of ester-linked ubiquitylation sites during TLR7 signalling increases the number of inter-ubiquitin linkages from 8 to 12 The Biochemical journal 479, 2419–2431 10.1042/BCJ20220510

20. Kelsall, I. R., Zhang, J., Knebel, A., Arthur, J. S. C., and Cohen, P. (2019) The E3 ligase HOIL-1 catalyses ester bond formation between ubiquitin and components of the Myddosome in mammalian cells Proc Natl Acad Sci U S A 116, 13293–13298 10.1073/pnas.1905873116

21. Otten, E. G., Werner, E., Crespillo-Casado, A., Boyle, K. B., Dharamdasani, V., Pathe, C., Santhanam, B., and Randow, F. (2021) Ubiquitylation of lipopolysaccharide by RNF213 during bacterial infection Nature 594, 111–116 10.1038/s41586-021-03566-4

22. Sakamaki, J. I., Ode, K. L., Kurikawa, Y., Ueda, H. R., Yamamoto, H., and Mizushima, N. (2022) Ubiquitination of phosphatidylethanolamine in organellar membranes Molecular cell 82, 3677–3692 e3611 10.1016/j.molcel.2022.08.008

23. Zhu, K., Suskiewicz, M. J., Hlousek-Kasun, A., Meudal, H., Mikoc, A., Aucagne, V., Ahel, D., and Ahel, I. (2022) DELTEX E3 ligases ubiquitylate ADP-ribosyl modification on protein substrates, Sci Adv 8, eadd4253 10.1126/sciadv.add4253

24. Zhu, K., Suskiewicz, M. J., Chatrin, C., Stromland, O., Dorsey, B. W., Aucagne, V., Ahel, D., and Ahel, I. (2023) DELTEX E3 ligases ubiquitylate ADP-ribosyl modification on nucleic acids, Nucleic acids research 10.1093/nar/gkad1119

25. Kelsall, I. R., McCrory, E. H., Xu, Y., Scudamore, C. L., Nanda, S. K., Mancebo-Gamella, P., Wood, N. T., Knebel, A., Matthews, S. J., and Cohen, P. (2022) HOIL-1 ubiquitin ligase activity targets unbranched glucosaccharides and is required to prevent polyglucosan accumulation, The EMBO journal 41, e109700 10.15252/embj.2021109700

26. Dearlove, E. L., Chatrin, C., Buetow, L., Ahmed, S. F., Schmidt, T., Bushell, M., Smith, B. O., and Huang, D. T. (2024) DTX3L ubiquitin ligase ubiquitinates single-stranded nucleic acids, bioRxiv 2024.2004.2002.587769 10.1101/2024.04.02.587769

27. Li, W., Garcia-Rivera, E. M., Mitchell, D. C., Chick, J. M., Maetani, M., Knapp, J. M., Matthews, G. M., Shirasaki, R., Simoes, R. d. M., Viswanathan, V., Pulice, J. L., Rees, M. G., Roth, J. A., Gygi, S. P., Mitsiades, C. S., Kadoch, C., Schreiber, S. L., and Ostrem, J. M. L. (2024) Highly specific intracellular ubiquitination of a small molecule, bioRxiv 2024.2001.2026.577493 10.1101/2024.01.26.577493

28. Wu, Q., Koliopoulos, M. G., Rittinger, K., and Stieglitz, B. (2022) Structural basis for ubiquitylation by HOIL-1, Front Mol Biosci 9, 1098144 10.3389/fmolb.2022.1098144

29. Wang, X. S., Cotton, T. R., Trevelyan, S. J., Richardson, L. W., Lee, W. T., Silke, J., and Lechtenberg, B. C. (2023) The unifying catalytic mechanism of the RING-between-RING E3 ubiquitin ligase family, Nat Commun 14, 168 10.1038/s41467-023-35871-z

30. Xu, X., Wang, Y., Zhang, Y., Wang, Y., Yin, Y., Peng, C., Gong, X., Li, M., Zhang, Y., Zhang, M., Tang, Y., Zhou, X., Liu, H., and Pan, L. (2023) Mechanistic insights into the enzymatic activity of E3 ligase HOIL-1L and its regulation by the linear ubiquitin chain binding, Sci Adv 9, eadi4599 10.1126/sciadv.adi4599

31. Matsumoto, M. L., Dong, K. C., Yu, C., Phu, L., Gao, X., Hannoush, R. N., Hymowitz, S. G., Kirkpatrick, D. S., Dixit, V. M., and Kelley, R. F. (2012) Engineering and structural characterization of a linear polyubiquitin-specific antibody, J Mol Biol 418, 134–144 10.1016/j.jmb.2011.12.053

32. Matsumoto, M. L., Wickliffe, K. E., Dong, K. C., Yu, C., Bosanac, I., Bustos, D., Phu, L., Kirkpatrick, D. S., Hymowitz, S. G., Rape, M., Kelley, R. F., and Dixit, V. M. (2010) K11-linked polyubiquitination in cell cycle control revealed by a K11 linkage-specific antibody, Molecular cell 39, 477–484 10.1016/j.molcel.2010.07.001

33. Newton, K., Matsumoto, M. L., Wertz, I. E., Kirkpatrick, D. S., Lill, J. R., Tan, J., Dugger, D., Gordon, N., Sidhu, S. S., Fellouse, F. A., Komuves, L., French, D. M., Ferrando, R. E., Lam, C., Compaan, D., Yu, C., Bosanac, I., Hymowitz, S. G., Kelley, R. F., and Dixit, V. M. (2008) Ubiquitin chain editing revealed by polyubiquitin linkage-specific antibodies, Cell 134, 668–678 10.1016/j.cell.2008.07.039

34. Michel, M. A., Swatek, K. N., Hospenthal, M. K., and Komander, D. (2017) Ubiquitin Linkage-Specific Affimers Reveal Insights into K6-Linked Ubiquitin Signaling, Molecular cell 10.1016/j.molcel.2017.08.020

35. Xu, G., Paige, J. S., and Jaffrey, S. R. (2010) Global analysis of lysine ubiquitination by ubiquitin remnant immunoaffinity profiling, Nat Biotechnol 28, 868–873 10.1038/nbt.1654

36. Akimov, V., Barrio-Hernandez, I., Hansen, S. V. F., Hallenborg, P., Pedersen, A. K., Bekker-Jensen, D. B., Puglia, M., Christensen, S. D. K., Vanselow, J. T., Nielsen, M. M., Kratchmarova, I., Kelstrup, C. D., Olsen, J. V., and Blagoev, B. (2018) UbiSite approach for comprehensive mapping of lysine and N-terminal ubiquitination sites, Nature structural & molecular biology 25, 631–640 10.1038/s41594-018-0084-y

37. Meyer, H. J., and Rape, M. (2014) Enhanced protein degradation by branched ubiquitin chains, Cell 157, 910–921 10.1016/j.cell.2014.03.037

38. Kirkpatrick, D. S., Hathaway, N. A., Hanna, J., Elsasser, S., Rush, J., Finley, D., King, R. W., and Gygi, S. P. (2006) Quantitative analysis of in vitro ubiquitinated cyclin B1 reveals complex chain topology, Nat Cell Biol 8, 700–710 10.1038/ncb1436

39. Valkevich, E. M., Sanchez, N. A., Ge, Y., and Strieter, E. R. (2014) Middle-down mass spectrometry enables characterization of branched ubiquitin chains, Biochemistry 53, 4979–4989 10.1021/bi5006305

40. Swatek, K. N., Usher, J. L., Kueck, A. F., Gladkova, C., Mevissen, T. E. T., Pruneda, J. N., Skern, T., and Komander, D. (2019) Insights into ubiquitin chain architecture using Ub-clipping, Nature 572, 533–537 10.1038/s41586-019-1482-y

41. Lechtenberg, B. C., and Komander, D. (2024) Just how big is the ubiquitin system? Nature structural & molecular biology 31, 210–213 10.1038/s41594-023-01208-z

42. Wenzel, D. M., Lissounov, A., Brzovic, P. S., and Klevit, R. E. (2011) UBCH7 reactivity profile reveals parkin and HHARI to be RING/HECT hybrids, Nature 474, 105–108 10.1038/nature09966

43. Cotton, T. R., and Lechtenberg, B. C. (2020) Chain reactions: molecular mechanisms of RBR ubiquitin ligases, Biochem Soc Trans 48, 1737–1750 10.1042/BST20200237

44. Walden, H., and Rittinger, K. (2018) RBR ligase-mediated ubiquitin transfer: a tale with many twists and turns, Nature structural & molecular biology 10.1038/s41594-018-0063-3

45. Sakamaki, J. I., and Mizushima, N. (2023) Ubiquitination of non-protein substrates, Trends Cell Biol 33, 991–1003 10.1016/j.tcb.2023.03.014

46. Hospenthal, M. K., Mevissen, T. E., and Komander, D. (2015) Deubiquitinase-based analysis of ubiquitin chain architecture using Ubiquitin Chain Restriction (UbiCRest), Nat Protoc 10, 349–361 10.1038/nprot.2015.018

47. De Cesare, V., Carbajo Lopez, D., Mabbitt, P. D., Fletcher, A. J., Soetens, M., Antico, O., Wood, N. T., and Virdee, S. (2021) Deubiquitinating enzyme amino acid profiling reveals a class of ubiquitin esterases, Proc Natl Acad Sci U S A 118, 10.1073/pnas.2006947118

48. Wegmann, S., Meister, C., Renz, C., Yakoub, G., Wollscheid, H. P., Takahashi, D. T., Mikicic, I., Beli, P., and Ulrich, H. D. (2022) Linkage reprogramming by tailor-made E3s reveals polyubiquitin chain requirements in DNA-damage bypass, Molecular cell 82, 1589–1602 e1585 10.1016/j.molcel.2022.02.016

49. Kelsall, I. R., McCrory, E. H., Xu, Y., Scudamore, C. L., Nanda, S. K., Mancebo-Gamella, P., Wood, N. T., Knebel, A., Matthews, S. J., and Cohen, P. (2021) HOIL-1-catalysed ubiquitylation of unbranched glucosaccharides and its activation by ubiquitin oligomers, bioRxiv 2021.2009.2010.459791 10.1101/2021.09.10.459791

50. Szczesna, M., Huang, Y., Lacoursiere, R. E., Bonini, F., Pol, V., Koc, F., Ward, B., Geurink, P. P., Pruneda, J. N., and Thurston, T. L. M. (2024) Bacterial esterases reverse lipopolysaccharide ubiquitylation to block host immunity, Cell Host Microbe 32, 913–924 e917 10.1016/j.chom.2024.04.012

51. Cotton, T. R., Cobbold, S. A., Bernardini, J. P., Richardson, L. W., Wang, X. S., and Lechtenberg, B. C. (2022) Structural basis of K63-ubiquitin chain formation by the Gordon-Holmes syndrome RBR E3 ubiquitin ligase RNF216, Molecular cell 82, 598–615 10.1016/j.molcel.2021.12.005

52. Lechtenberg, B. C., Rajput, A., Sanishvili, R., Dobaczewska, M. K., Ware, C. F., Mace, P. D., and Riedl, S. J. (2016) Structure of a HOIP/E2∼ubiquitin complex reveals RBR E3 ligase mechanism and regulation, Nature 529, 546–550 10.1038/nature16511

53. Corcilius, L., and Payne, R. J. (2013) Stereoselective synthesis of sialylated tumor-associated glycosylamino acids, Org Lett 15, 5794–5797 10.1021/ol402845e

54. Michel, M. A., Komander, D., and Elliott, P. R. (2018) Enzymatic Assembly of Ubiquitin Chains Methods, Mol Biol 1844, 73–84 10.1007/978-1-4939-8706-1_6

55. Signor, L., and Boeri Erba, E. (2013) Matrix-assisted laser desorption/ionization time of flight (MALDI-TOF) mass spectrometric analysis of intact proteins larger than 100 kDa, J Vis Exp 10.3791/50635

56. Aragao, D., Aishima, J., Cherukuvada, H., Clarken, R., Clift, M., Cowieson, N. P., Ericsson, D. J., Gee, C. L., Macedo, S., Mudie, N., Panjikar, S., Price, J. R., Riboldi-Tunnicliffe, A., Rostan, R., Williamson, R., and Caradoc-Davies, T. T. (2018) MX2: a high-flux undulator microfocus beamline serving both the chemical and macromolecular crystallography communities at the Australian Synchrotron, Journal of synchrotron radiation 25, 885–891 10.1107/S1600577518003120

57. Kabsch, W. (2010) Xds Acta crystallographica Section D, Biological crystallography 66, 125–132 10.1107/S0907444909047337

58. Evans, P. R., and Murshudov, G. N. (2013) How good are my data and what is the resolution?, Acta crystallographica Section D, Biological crystallography 69, 1204–1214 10.1107/S0907444913000061

59. McCoy, A. J., Grosse-Kunstleve, R. W., Adams, P. D., Winn, M. D., Storoni, L. C., and Read, R. J. (2007) Phaser crystallographic software, J Appl Crystallogr 40, 658–674 10.1107/S0021889807021206

60. Vijay-Kumar, S., Bugg, C. E., and Cook, W. J. (1987) Structure of ubiquitin refined at 1.8 A resolution, J Mol Biol 194, 531–544 10.1016/0022-2836(87)90679-6

61. Emsley, P., Lohkamp, B., Scott, W. G., and Cowtan, K. (2010) Features and development of Coot, Acta crystallographica Section D, Biological crystallography 66, 486–501 10.1107/S0907444910007493

62. Afonine, P. V., Moriarty, N. W., Mustyakimov, M., Sobolev, O. V., Terwilliger, T. C., Turk, D., Urzhumtsev, A., and Adams, P. D. (2015) FEM: feature-enhanced map, Acta crystallographica Section D, Biological crystallography 71, 646–666 10.1107/S1399004714028132

63. Liebschner, D., Afonine, P. V., Baker, M. L., Bunkoczi, G., Chen, V. B., Croll, T. I., Hintze, B., Hung, L. W., Jain, S., McCoy, A. J., Moriarty, N. W., Oeffner, R. D., Poon, B. K., Prisant, M. G., Read, R. J., Richardson, J. S., Richardson, D. C., Sammito, M. D., Sobolev, O. V., Stockwell, D. H., Terwilliger, T. C., Urzhumtsev, A. G., Videau, L. L., Williams, C. J., and Adams, P. D. (2019) Macromolecular structure determination using X-rays, neutrons and electrons: recent developments in Phenix, Acta Crystallogr D Struct Biol 75, 861–877 10.1107/S2059798319011471

64. Pettersen, E. F., Goddard, T. D., Huang, C. C., Meng, E. C., Couch, G. S., Croll, T. I., Morris, J. H., and Ferrin, T. E. (2021) UCSF ChimeraX: Structure visualization for researchers, educators, and developers, Protein Sci 30, 70–82 10.1002/pro.3943

65. Geoghegan, K. F., Dixon, H. B., Rosner, P. J., Hoth, L. R., Lanzetti, A. J., Borzilleri, K. A., Marr, E. S., Pezzullo, L. H., Martin, L. B., LeMotte, P. K., McColl, A. S., Kamath, A. V., and Stroh, J. G. (1999) Spontaneous alpha-N-6-phosphogluconoylation of a “His tag” in Escherichia coli: the cause of extra mass of 258 or 178 Da in fusion proteins, Analytical biochemistry 267, 169–184 10.1006/abio.1998.2990

